# Human Precision-Cut Kidney Slices as a Translational Platform for Modelling Fibrotic, Inflammatory, Metabolic and Drug-Induced Kidney Injury

**DOI:** 10.64898/2026.11.25.768989

**Authors:** Giuseppe Abbate, Laura Sabater, Lee Reed, Hannah Paish, Charlotte Bragg, Nicola Griffiths, Rebecca Scott, Sarah Worthington, Eleni Tzortzopoulou, Luke Patterson, Ben Barksby, Rachel Burgoyne, Lucy Bates, Aleksandra Gruevska, Zoe Hall, Li Chen, Mathieu Petitjean, Neil S Sheerin, Fiona Oakley, Derek A Mann, Jelena Mann, Lee A Borthwick

## Abstract

**Introduction:** Chronic kidney disease (CKD) is characterised by progressive fibrosis and inflammation. Preclinical models of these processes translate poorly to human CKD. Animal models are limited by species differences, while two-dimensional cultures and organoids do not reproduce the three-dimensional, multicellular environment of the human kidney. We sought to develop and characterise precision-cut human kidney slices (PCKS) as an *ex-vivo* platform for studying renal disease and testing therapeutics.

**Methods:** PCKS were prepared from human donor kidneys not utilised for transplantation and cultured in a proprietary bioreactor for up to six days. Tissue integrity, fibrogenesis, inflammation, lipid handling, and drug-induced injury were assessed by histology, digital pathology, secreted protein quantification, mass spectrometry-based lipidomics, and bulk RNA sequencing, with and without pharmacological intervention.

**Results:** PCKS retained native cortical architecture and viability throughout culture. Slices underwent spontaneous, time-dependent fibrogenesis that was enhanced by transforming growth factor beta 1, abrogated by an ALK5 inhibitor, and attenuated by nintedanib but not pirfenidone. Single-fibre digital pathology resolved treatment-dependent changes in collagen content, morphology and architecture. Proinflammatory stimuli induced stimulus-selective cytokine and chemokine secretion through NF-κB, which was suppressed by an IKK2 inhibitor. Fatty acid treatment reproduced features of fatty kidney disease, including triacylglycerol accumulation and dose-dependent inflammation, while nephrotoxic drugs produced characteristic injury detectable across multiple longitudinal readouts.

**Conclusions:** Human PCKS maintain renal architecture and viability ex-vivo and reproduce fibrotic, inflammatory, metabolic, and toxic responses at transcriptomic, proteomic and lipidomic scales. PCKS provide a physiologically relevant, human-based platform for renal disease modeling and drug discovery.

**Translational Statement:** The development of effective therapies for chronic kidney disease is hindered by preclinical models that inadequately reproduce human renal biology. We demonstrate that human precision-cut kidney slices maintain native tissue architecture and support the modelling of fibrosis, inflammation, metabolic dysfunction and drug-induced injury. The platform captures disease-relevant molecular and tissue responses and distinguishes pharmacological effects, including inhibition of fibrogenesis and inflammatory signalling. By enabling therapeutic evaluation directly in human kidney tissue, this approach offers a translational bridge between conventional preclinical models and clinical research, with potential to improve drug discovery and reduce reliance on animal experimentation.

## Introduction

Kidney fibrosis is a hallmark of progressive chronic kidney disease (CKD) and represents a major challenge in nephrology research^1^. Physiologically relevant experimental models are critical for studying fibrotic mechanisms and evaluating potential therapies. While animal models and *in-vitro* cell culture systems have contributed significantly to our understanding of kidney fibrosis, they have limitations in fully recapitulating the complex cellular interactions and tissue-specific responses observed in the human kidney^2^. Precision-cut kidney slices have previously been used to model renal fibrosis in murine and human tissue^3,4^. Building on these previous studies, we have established a novel *ex vivo* human precision-cut kidney slice (PCKS) model that uses a bioreactor to preserve tissue viability, native three-dimensional architecture and the multicellular environment for six days, providing a more physiologically relevant system for studying kidney fibrosis.

PCKS offer several advantages over traditional experimental models, including the maintenance of the intricate spatial relationships between different cell types and the preservation of the extracellular matrix (ECM), which plays a crucial role in fibrosis. Moreover, PCKS provide a unique opportunity to investigate human-specific cellular and molecular mechanisms, which are often not well-conserved across species, thereby improving the translational potential of research findings. In addition, the use of human kidneys, which were not utilised for transplantation, offers an ethically sound and valuable source of tissue for research purposes.

Here, we describe the preparation, culture, and characterization of PCKS and their application as a model of acute and chronic kidney damage, inflammation and fibrogenesis. We discuss the advantages and limitations of the model and its potential to advance understanding of fibrotic and inflammatory mechanisms and support the development of novel therapies.

## Materials and methods

Detailed protocols, including reagent sources, concentrations and analytical parameters, are provided in the Supplementary Materials and Methods.

### Human Kidney Tissue

Kidneys from deceased UK donors were obtained through the Newcastle Institute of Transplantation Tissue Biobank. Research consent was obtained by Specialist Nurses in Organ Donation, under NHS Research Ethics Committee approval (REC 22/NE/0064).

### Precision-Cut Kidney Slice Culture

Precision-cut kidney slices (PCKS) were prepared from renal cortex using a vibrating microtome. They were cultured on Transwell inserts on a rocking bioreactor platform, as previously described.^5,6^ Two formats were used: 8 mm slices in 12-well plates, and 3 mm slices in a 96-well format for higher-throughput testing (Supplementary Figure S1). Following a 24 h rest period, media and treatments were refreshed daily, and supernatants were collected for analysis of secreted markers.

### Disease Models

Fibrosis was induced with TGF-β1, alone or with PDGF-ββ, for 72 h. This was done in the presence or absence of an ALK5 inhibitor, nintedanib or pirfenidone. Inflammation was induced by a single 24 h challenge with IL-1α, IL-1β or LPS, with or without IKK2 inhibition. Lipotoxicity was modelled by 72 h exposure to free fatty acids (0.25–2 mM). Acute nephrotoxic injury was modelled by 24 h exposure to cisplatin, gentamicin or staurosporine, and tissue was assessed at peak injury and after 24 h of recovery.

### Histology and Digital Pathology

Formalin-fixed sections were stained with H&E, PAS, Picrosirius Red and αSMA, and cryosections with Oil Red O. Fibrosis was quantified on whole-slide Picrosirius Red images using FibroNest™ (PharmaNest), an AI-based single-fibre image analysis platform. It measures collagen content, fibre morphometry and fibrosis architecture, and combines them into a Phenotypic Fibrosis Composite Score.^7^

### Biochemical and Viability Assays

Secreted markers of fibrosis, inflammation and injury were measured in culture supernatants by ELISA and by Meso Scale Discovery multiplex immunoassay. Tissue viability and injury were assessed by LDH release, ATP content, resazurin reduction, membrane integrity and caspase-3/7 activity.

### Transcriptomic and Lipidomic Profiling

Bulk RNA sequencing (stranded mRNA; >30 million reads per sample) was performed at the Newcastle University Genomics Core Facility. Differential expression was determined using DESeq2 (adjusted P<0.001, log₂ fold change >2), with pathway analysis in Ingenuity Pathway Analysis. Lipids were extracted by the Folch method and profiled by untargeted ultra-high-performance liquid chromatography–high-resolution mass spectrometry.

## Results

### Spontaneous fibrogenesis is a feature of human PCKS cultures and can be attenuated by therapeutic treatment with Nintedanib

We previously described a proprietary bioreactor and methodology for the optimal *ex-vivo* cultivation of precision-cut human liver slices^5^. With minor protocol modifications, we generated human Precision Cut Kidney Slices (PCKS). To generate PCKS, freshly harvested human kidney cortex tissue from n=42 individual donors (Table 1) was subjected to coring, agarose embedding and preparation of 3mm or 8-mm-diameter, 250-µm-thick PCKS (Supplementary Figure S1). Slices were cultured in our patented bioreactor for six days, with longitudinal sampling of conditioned media and tissue. These 42 donors represent the total tissue pool for the study; individual experiments used subsets of these donors, as indicated in the figure legends and methods.

**Table 1:** Baseline characteristics of declined donor organs used in this study (n=42)

| **Age** | **Sex** | **Reason declined** |
|---|---|---|
| **Age, median (range)** |  | 64.5 (20-79) |
| 63 | M | No reason listed |
| 51 | M | No reason listed |
| 65 | M | No reason listed |
| 72 | M | No reason listed |
| 65 | M | No reason listed |
| 53 | F | No reason listed |
| 57 | F | No reason listed |
| 73 | F | Damaged |
| 58 | F | Anatomical |
| 69 | F | Severe atheroma of aorta extending into left renal artery, stripped of intima |
| 76 | M | Benign cyst |
| 56 | F | No reason listed |
| 71 | F | No reason listed |
| 64 | M | Diseased patch in arteries, hematoma on kidney near upper pole |
| 69 | F | No reason listed |
| 71 | F | Poor perfusion of kidney, improved on extra perfusion |
| 63 | M | Poor perfusion of kidney |
| 68 | F | Anatomical |
| 75 | F | No reason listed |
| 63 | F | No reason listed |
| 70 | M | No reason listed |
| 55 | F | No reason listed |
| 75 | M | No reason listed |
| 68 | F | No reason listed |
| 79 | F | 1cm cyst, haemorrhage - small kidneys |
| 65 | M | Atherosclerotic patch, severe & mass 2cm in middle of kidney with cyst |
| 62 | M | Adherent perirenal fat |
| 58 | M | Severe adherant fat, large capsular tear - moderate damage |
| 58 | M | Densely adherant fat, not possible to fully excise |
| 64 | F | High BMI - +24h since retrieval |
| 65 | F | Nil arterial dissection |
| 44 | F | Anatomic - histopathology on liver |
| 72 | M | Anatomical - cyst |
| 73 | M | Anatomical/organ damage |
| 63 | M | Possible aneurysm - black blood pooled |
| 70 | F | L kidney with 1cm cyst - used the right one |
| 43 | M | No reason listed |
| 63 | M | Lesion on donor lung |
| 20 | M | Damaged |
| 58 | F | Poor perfusion and cholesterol emboli |
| 74 | F | Anatomical, NSR |
| 59 | M | Anatomy |
| **Sex (M/F)** |  | 21/21 |

*n = 42 donors. Age is presented as median (range) in years. Reasons for decline as documented at time of offer, where applicable.*

The first objective of our investigation involved the histological examination of the bioreactor-cultivated PCKS. By utilising haematoxylin and eosin staining, we found the slices demonstrated preserved structural integrity and closely resembled the typical morphology expected for human kidney cortex (Figure 1A), with mild features of tubular injury consistent with the donor ischaemic time typical of deceased-donor kidneys also noted in a subset of slices. Following this, we employed picrosirius red (PSR) and alpha smooth muscle actin (αSMA) immunohistochemical staining to assess fibrosis and visualise myofibroblasts. These parameters were analysed both at the onset (0hrs) and over the duration of the culture period (up to 144hrs) (Figure 1A). The observations revealed a time-dependent, spontaneous (i.e. exogenous-stimulus-independent) fibrotic progression in the tissue. This progression may reflect, at least in part, a fibrotic response to the warm and cold ischaemic injury sustained by deceased-donor kidneys prior to retrieval – a phenomenon well documented following pre-renal acute kidney injury – and is itself of interest as a model of post-ischaemic renal fibrosis; this progression was not accompanied by an increase in numbers of αSMA-positive cells within the same timeframe. This suggests that collagen deposition in this model may reflect activation of resident stromal cells and enhanced extracellular matrix production rather than expansion or overt differentiation of αSMA-positive myofibroblasts. Concurrently, longitudinal evaluation of the conditioned media derived from the PCKS highlighted an autonomous increment in the secretion of Collagen1α1 and TIMP-1 up to 96hrs, thereby providing further corroboration of the evolving structural modifications within the slices in absence of external stimulation (Figure 1B). Conversely, secretion of hyaluronic acid and the pro-inflammatory marker MCP-1 remained elevated but unchanged during culture (Figure 1B).

**Figure 1.**
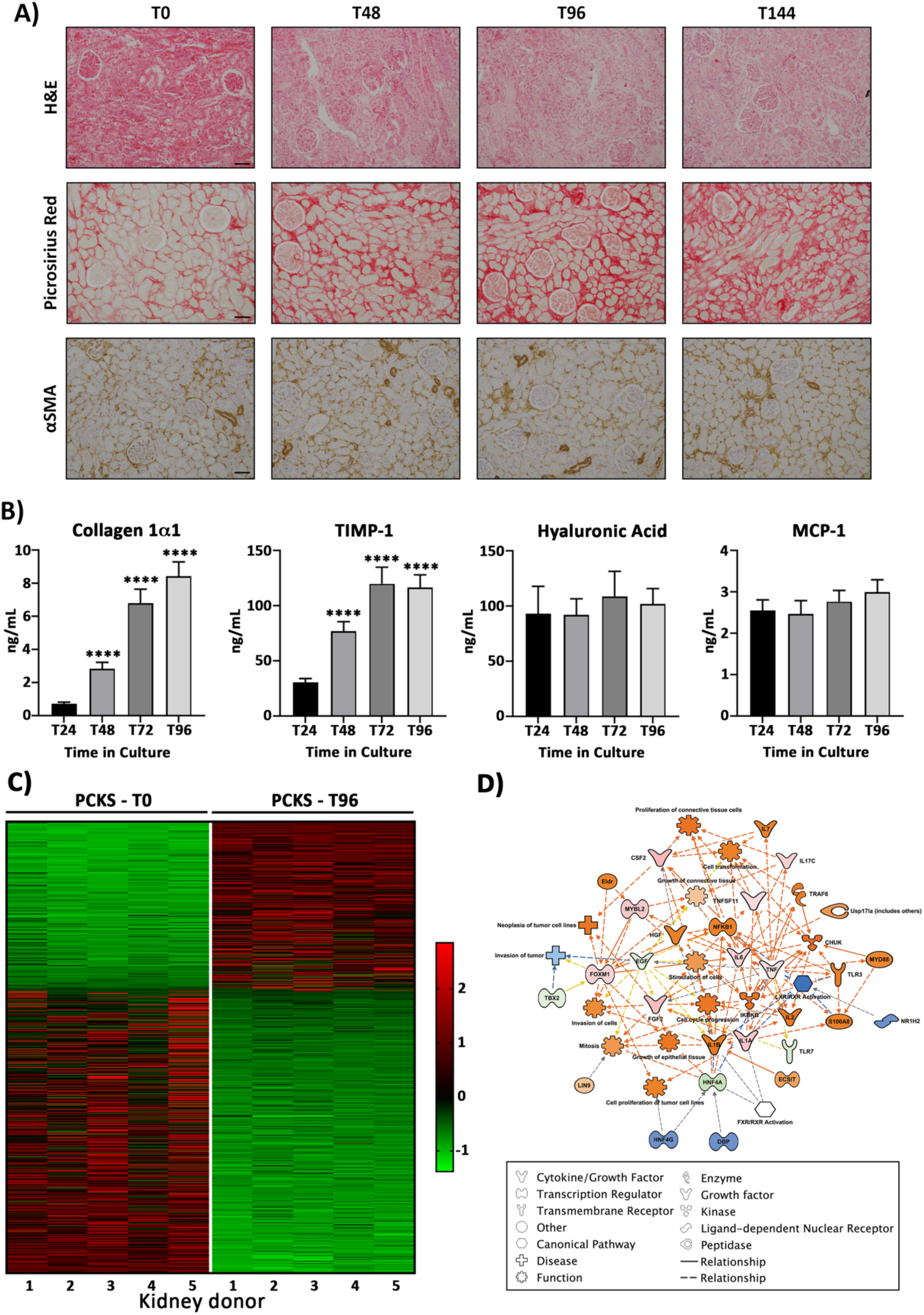
Bioreactor-cultured PCKS maintain viability for at least 6 days in culture. (A) Precision-cut kidney slices (PCKS) were prepared from unused donor kidneys and immediately fixed (T0) or fixed after 48hrs (T48), 96hrs (T96) or 144hrs (T144) of bioreactor culture. Representative ×100 magnification images of H&E, picrosirius red and α-smooth muscle actin (αSMA) staining demonstrate maintenance of tissue integrity. Scale bars = 100 µm. (B) Quantified levels of collagen 1α1, TIMP-1, hyaluronic acid and MCP-1 in PCKS culture media after 24hrs (T24), 48hrs (T48), 72hrs (T72) and 96hrs (T96). Data are mean ± SEM (n=35 donors for collagen 1α1; n=17 for TIMP-1; n=6 for hyaluronic acid; n=15 for MCP-1), analysed by one-way ANOVA with Dunnett’s multiple comparisons test versus T24; ****p<0.0001. (C) Heat map of n=2497 differentially expressed genes (log2 fold change, p<0.001) between T0 and T96 PCKS (n=5 donors). (D) Ingenuity Pathway Analysis of genes differentially expressed during PCKS culture and the pathways regulated.

### Confirming PCKS sensitivity to drug-induced acute renal damage

Before applying PCKS to model the fibrotic, inflammatory and metabolic disease processes described below, we first confirmed that the platform retains the ability to detect drug-induced nephrotoxicity. PCKS were exposed to two clinically relevant nephrotoxic agents – cisplatin (150µM) and gentamicin (20µg/mL), together with staurosporine (10µM, used as a positive control for apoptosis) – for 24 or 48hrs. Histological assessment by Periodic Acid-Schiff (PAS) staining confirmed that all three agents induced features consistent with clinically observed drug-induced nephrotoxicity, including loss of proximal tubular brush border integrity and tubular epithelial vacuolisation (Supplementary Figure S2A). Drug exposure induced a significant increase in Caspase 3/7 activity, most notably with staurosporine at 24hrs, alongside a reduction in metabolic activity (Cell Titre-Glo) that reached significance for all three agents by 48hrs (Supplementary Figure S2B). A staurosporine dose–response confirmed concentration-dependent loss of metabolic activity (Supplementary Figure S2C). These data confirm that PCKS can detect and discriminate compound-induced nephrotoxic effects, supporting their use as a platform for early drug safety screening.

To identify the molecular events underpinning spontaneous fibrogenesis, we performed bulk RNA sequencing of PCKS from five individual donor experiments. A comparative analysis was performed between the initial time point (0hrs), and the time point that registered the most significant changes in the longitudinally collected soluble markers (96hrs), discovering 2497 differentially expressed genes (Figure 1C, Supplementary Table 1). Ingenuity Pathway Analysis (IPA) revealed changes in pathways implicated in growth of connective tissue, stimulation of cells, cell cycle progression and growth of epithelial cells with evidence of alterations in NF-κB signalling (NFKB1 and IKBKB), inflammatory signalling molecules (IL6, IL1A, IL1B and TNF) and growth factors (HGF, FGF2 and EGF) (Figure 1D). Complementary Hallmark, KEGG and Gene Ontology enrichment analyses confirmed activation of profibrotic, inflammatory and proliferative programmes alongside alterations in tubular epithelial structure and renal transport pathways (Supplementary Figure S3).

To determine if the PCKS platform can be used to therapeutically monitor the effects of known anti-fibrotic molecules we exposed cultures to either Pirfenidone or Nintedanib in the absence of exogenous challenge to determine effects on secretion of fibrogenic proteins in the culture media. Levels of Collagen 1α1 in PCKS media again demonstrated a spontaneous increase at 72hrs and 96hrs of culture and this was attenuated by exposure to Nintedanib. However, Collagen 1α1 was unaffected by treatment with Pirfenidone (Figure 2A). Similarly, Nintedanib also attenuated secretion of other fibrogenic proteins including fibronectin, IL-6 and TIMP-1, but again we did not observe inhibitory effects from Pirfenidone treatment (Figure 2B-D). Hence, PCKS can discriminate between anti-fibrotic molecules, although the lack of a Pirfenidone effect, despite its clinical use in Idiopathic Pulmonary Fibrosis (IPF), may reflect the single dose tested rather than optimal therapeutic exposure. To corroborate these data, we carried out bulk RNAseq analysis on the PCKS collected at the end of culture period. Heat map analysis clearly shows a powerful modulatory effect of Nintedanib on gene expression which was not observed with Pirfenidone (Figure 2E, Supplementary Table 2). Complementary KEGG, Reactome and Gene Ontology enrichment analyses of these nintedanib-modulated genes confirmed attenuation of a co-ordinated collagen and extracellular matrix remodelling, inflammatory and metabolic programme (Supplementary Figure S4).

**Figure 2.**
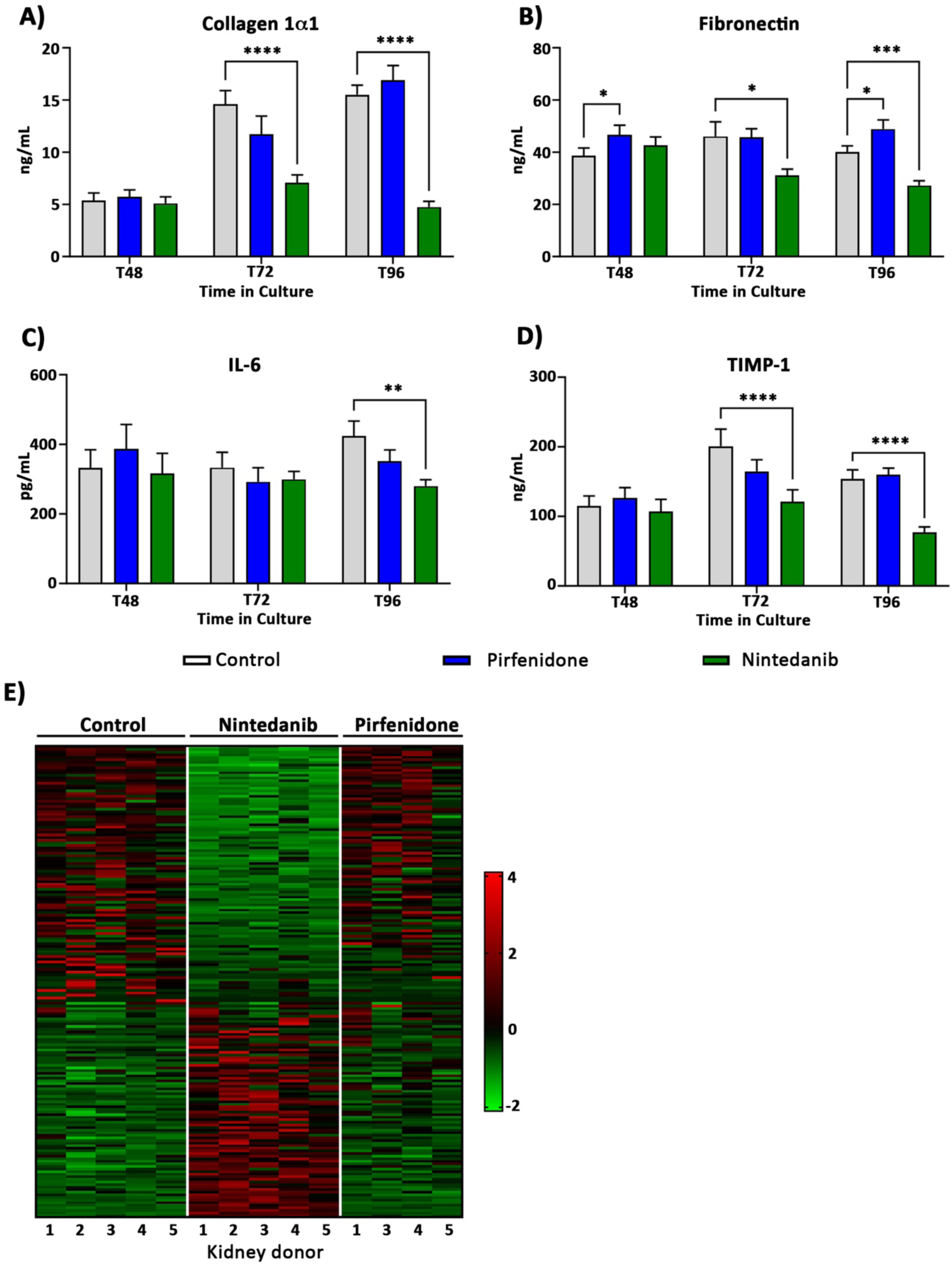
Testing clinically approved anti-fibrotics in human PCKS. (A–D) Quantified levels of collagen 1α1, fibronectin, IL-6 and TIMP-1 in PCKS culture media after 48hrs (T48), 72hrs (T72) and 96hrs (T96) in the control group and in the presence of pirfenidone or nintedanib. Data are mean ± SEM (n=6 donors), analysed by one-way ANOVA with Dunnett’s multiple comparisons test versus control; ****p<0.0001. (E) Heat map of n=182 differentially expressed genes (log2 fold change, p<0.05) between control, nintedanib and pirfenidone groups at T96 (n=5 donors).

### Exogenous TGF-β1 enhances spontaneous PCKS fibrogenesis

TGF-β1 is a potent profibrogenic cytokine that promotes fibrosis *via* well characterised paracrine and autocrine mechanisms.^8^ We therefore asked if the spontaneous fibrogenesis observed in human PCKS can be accentuated by exposing PCKS cultures to exogenously applied TGF-β1. We additionally asked if blocking TGFβR1 signalling with an ALK5 inhibitor (ALK5i) can prevent spontaneous fibrogenesis and TGF-β1 induced fibrogenesis by quantifying secretion of Collagen1α1, Fibronectin, IL-6 and TIMP-1 proteins in the PCKS culture media as fibrogenic biomarker readouts (Figure 3A-D). All four proteins displayed a robust and significant dynamic increase in secretion in response to challenge with TGF-β1, an effect that was completely abrogated by ALK5i. Similarly, the spontaneous fibrogenesis observed in non-challenged PCKS was also attenuated by ALK5i, indicating a contribution of TGF-β1/TGF-βR1 signalling in the spontaneous fibrogenesis observed in PCKS. At the end of the culture period, extracted RNA from PCKS was sequenced, which upon transcript analysis showed significant changes in the expression of 106 genes (Figure 3E, Supplementary Table 3). Complementary Reactome and Gene Ontology enrichment analyses confirmed that TGF-β1-regulated genes were enriched for extracellular matrix organisation, collagen formation, matrix degradation, growth factor activity and cytokine activity, supporting activation of a focused profibrotic transcriptional programme in human PCKS (Supplementary Figure S5).

**Figure 3.**
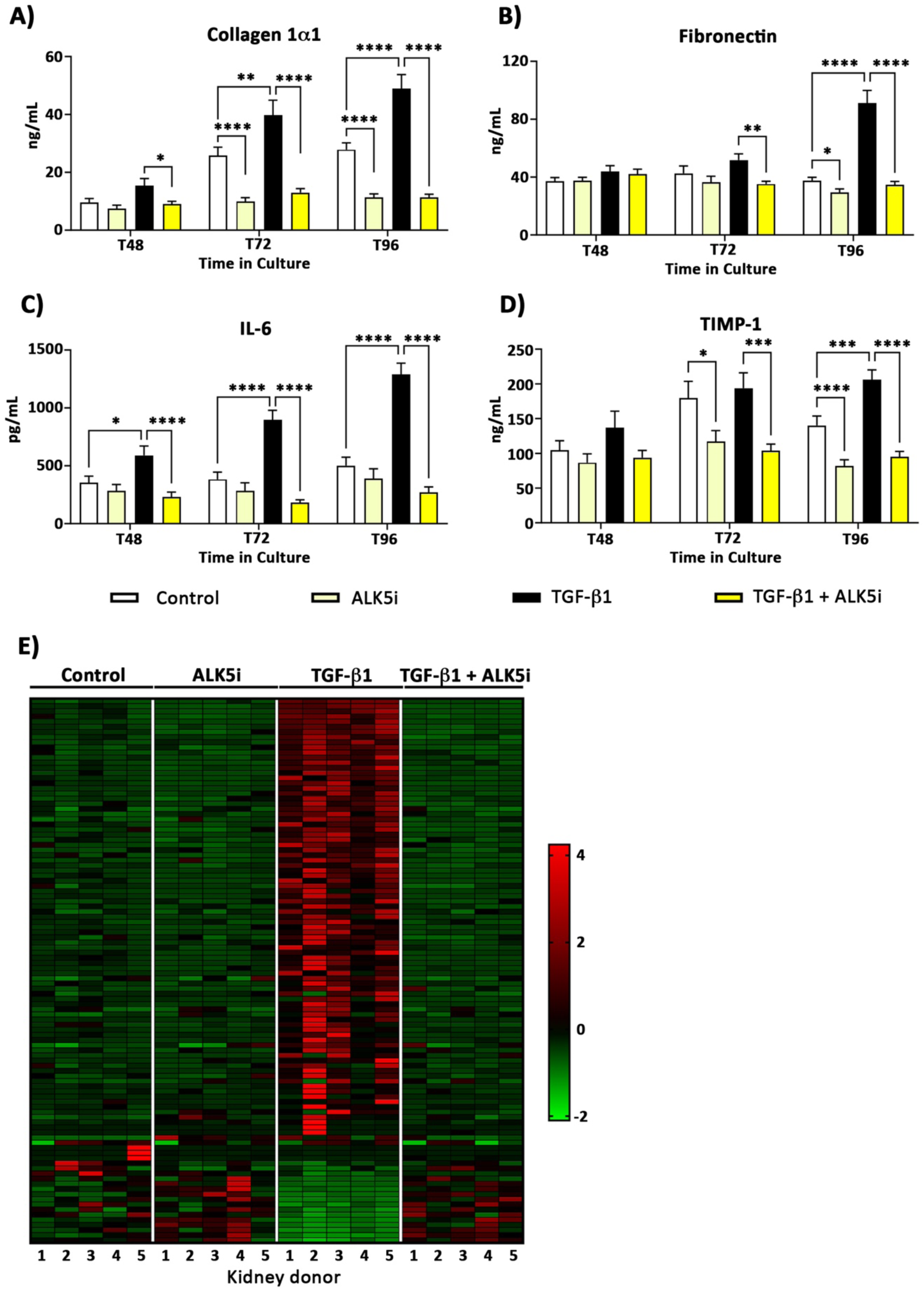
Inducing fibrosis with the pro-fibrogenic stimulus TGF-β1 in human PCKS. (A–D) Quantified levels of collagen 1α1, fibronectin, IL-6 and TIMP-1 in PCKS culture media after 48hrs (T48), 72hrs (T72) and 96hrs (T96) in the control group and in the presence of ALK5i, TGF-β1, or TGF-β1 + ALK5i. Data are mean ± SEM (n=7 donors), analysed by two-way ANOVA with Dunnett’s multiple comparisons; ****p<0.0001. (E) Heat map of n=106 differentially expressed genes between control, ALK5i, TGF-β1 and TGF-β1 + ALK5i groups at T96 (n=5 donors).

Having established that PCKS undergo extensive fibrotic remodelling in response to TGF-β1 we were interested to investigate the molecular architecture of the ECM deposited into the cultured kidney tissue. To this end in a further PCKS experiment, we spatially imaged and quantified deposited collagen I/III protein within control, TGF-β1 and TGF-β1+ Alk5i treated PCKS using the FibroNest^TM^ digital Pathology method ^7,9^. This high-resolution, single fiber quantitative image analysis method quantifies the phenotype of fibrosis in 313 dimensions including collagen deposition (12 parameters, including Collagen Area ratio), fiber morphometry (252 parameters) and fibrosis architecture (49 parameters). The traits that best captured the remodeling of fibrosis between the control and TGF-β1 groups were selected and assembled into equally weighted, normalized composite scores. These scores characterize the morphometry, collagen content, and architectural phenotypes of fibrosis. The resulting phenotypic composite score effectively recapitulates the overall fibrotic phenotype. Comparison of picrosirius red stained tissues (Figure 4A, top row) with images processed by FibroNest^TM^ for collagen architecture mapping (Fig 4A, second row), collagen density (Figure 4A, third row) and collagen assembly (Fig 4A, bottom row), demonstrate the investigative power of the technology when combined with PCKS. Quantification of these parameters (Figure 4B-D) revealed that TGF-β1 promotes an increase in total collagen I/III in the tissue primarily through an accumulation of assembled (complex skeleton^7^) collagen fibres and a decrease in fine (simple skeleton^7^) collagen fibres, indicative of active fibrogenesis^10^, with this function being dependent on canonical TGFβR1 signalling as all quantified parameters were effectively suppressed by ALK5i (Figure 4B). Furthermore, morphological assessment of the collagen fibres using the FibroNest Fibrosis Morphometric Score, confirmed that TGF-β1 induced phenotypic and architectural changes to the collagen fibres in the PCKS, and this was suppressed by ALK5i (Figure 4C-D). These investigations highlight the advantages of PCKS modelling for detailed multi-dimensional molecular analysis of the fibrogenic process and deeper interrogation of matrix deposition and architecture.

**Figure 4.**
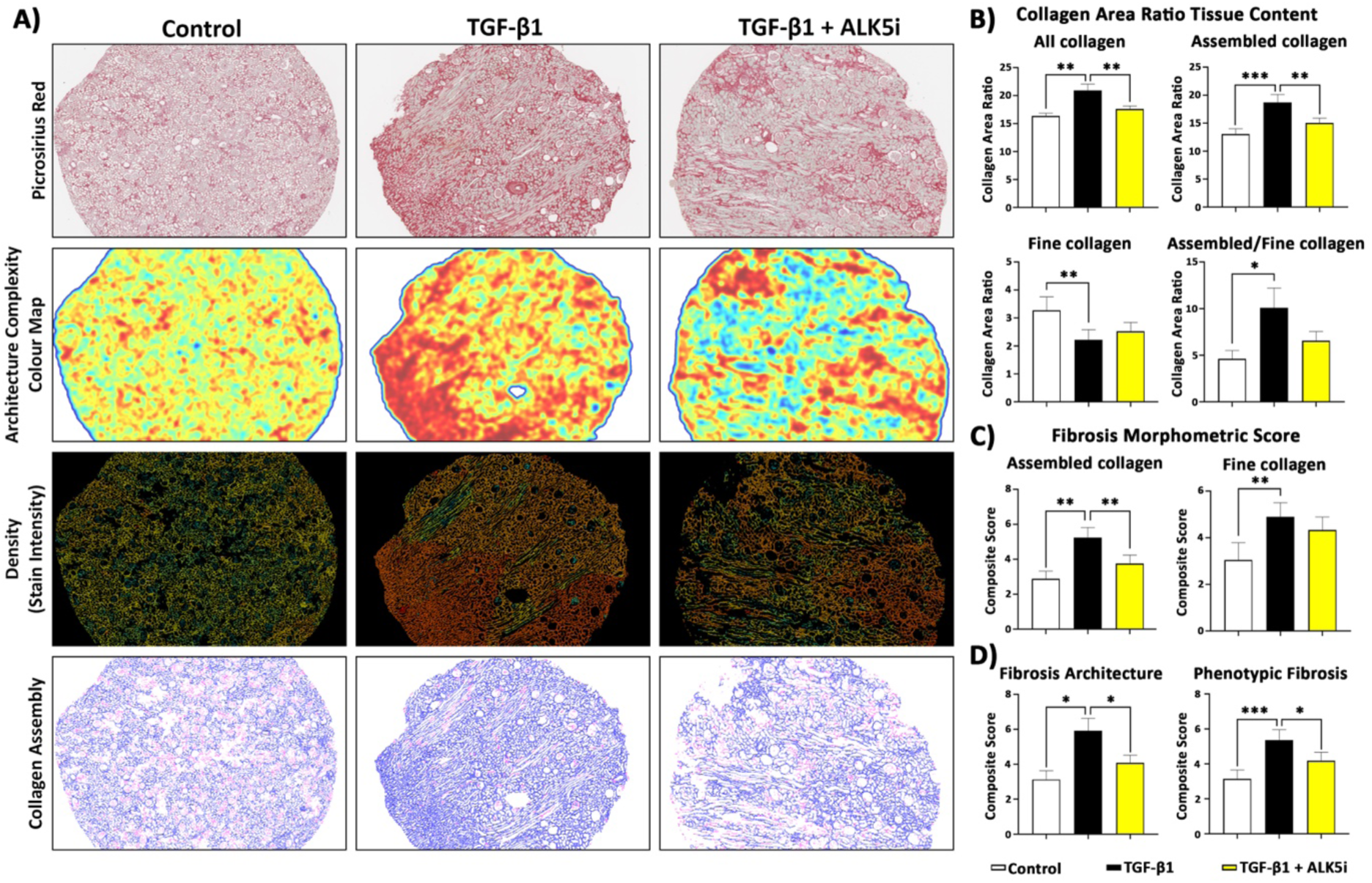
Using digital pathology (FibroNest™) to evaluate collagen architecture, density and assembly in PCKS. PCKS were fixed after 96hrs of bioreactor culture for control, TGF-β1 and TGF-β1 + ALK5i groups. (A) Representative ×20 magnification images show picrosirius red staining (top row) and FibroNest™ outputs for collagen architecture colour map, collagen density and collagen assembly. (B) Collagen area ratio relative to tissue content for total, assembled and fine collagen, and the assembled/fine ratio. (C–D) Fibrosis Morphometric Composite Scores for assembled collagen, fine collagen, fibrosis architecture and phenotypic fibrosis. Data are mean ± SEM (n=5 donors), analysed by two-way ANOVA with Dunnett’s multiple comparisons; ****p<0.0001.

We next investigated whether TGF-β1 induced fibrogenesis in PCKS could be attenuated by Pirfenidone or Nintedanib treatment. Similarly to the spontaneous model, only Nintedanib treatment of PCKS attenuated fibrogenic protein secretion (Fig 5A-D) and changes in gene expression (Fig 5E, Supplementary Table 4). Complementary KEGG, Reactome and Gene Ontology enrichment analyses of these nintedanib-modulated genes confirmed attenuation of a coordinated collagen and extracellular matrix remodelling and inflammatory programme in TGF-β1-stimulated PCKS (Supplementary Figure S6).

**Figure 5.**
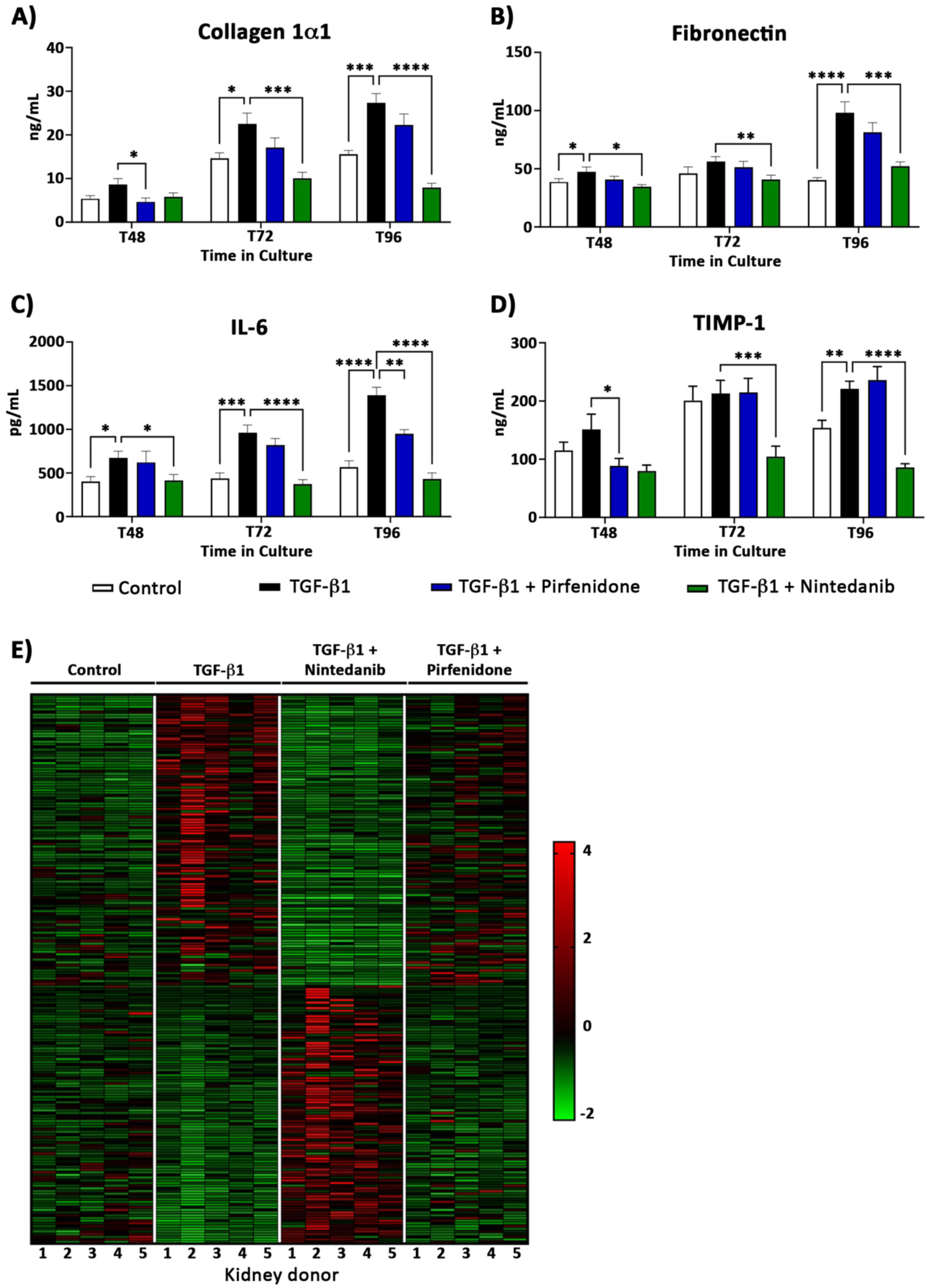
Testing clinically approved anti-fibrotics in human PCKS in combination with pro-fibrotic TGFβ1 stimulus. (A–D) Quantified levels of collagen 1α1, fibronectin, IL-6 and TIMP-1 in PCKS culture media after 48hrs (T48), 72hrs (T72) and 96hrs (T96) in the control group and in the presence of TGFβ1, TGFβ1 + nintedanib or TGFβ1 + pirfenidone. Data are mean ± SEM (n=6 donors), analysed by two-way ANOVA with Dunnett’s multiple comparisons; ****p<0.0001. (E) Heat map of n=204 differentially expressed genes between control, TGF-β1, TGF-β1 + nintedanib and TGF-β1 + pirfenidone groups at T96 (n=5 donors).

PCKS were also treated with a combination of TGFβ1 and a second well established profibrogenic molecule, PDGFβ^5,11^. We again observed elevated expression of secreted fibrogenic proteins (Figure S7A–D), with these effects being abrogated by ALK5i, indicating that TGFβ1 was the dominant profibrogenic factor in these experiments. Bulk RNAseq on RNA isolated from PCKS at the termination of the experiment (96hrs) documented that the expression of 334 transcripts was altered in response to combined TGF-β1/PDGFββ treatment (Figure S7E, Supplementary Table 5). As previously shown for spontaneous fibrogenesis (Figure 2), fibrogenesis induced by combined TGFβ1/PDGFββ treatment of PCKS was robustly inhibited by nintedanib, whereas pirfenidone broadly lacked efficacy in this model (Figure S8A–E, Supplementary Table 6).

### Human PCKS are responsive to inflammatory stimuli via activation of NF-κB

We next transitioned from the exploration of fibrogenesis and fibrosis-related changes in PCKS to assessing the PCKS as a platform for studying renal inflammation. To do this, we challenged PCKS to a range of proinflammatory stimuli, specifically, Interleukin 1 Beta (IL-1β), Interleukin 1 Alpha (IL-1α) and Lipopolysaccharide (LPS), for 24hrs and assessed the secretion of 111 cytokines and chemokines in the conditioned media using a Multiplex ELISA (MSD). For all stimuli tested, there was a statistically significant increase in the secretion of numerous chemokines and cytokines in response to *ex-vivo* challenge. PCKS challenged with IL-1β upregulated secretion of n=22 markers (Figure 6A) whereas challenge with IL-1α and LPS increased secretion of n=21 (Figure S9A–B) and n=26 (Figure S9C–D) markers, respectively. Comparing proteins induced in response to the three different stimuli, 12 inflammatory mediators (GRO-α, IL-6, M-CSF, Fractalkine, MCP-1, MCP-2, G-CSF, MCP-4, IL-7, TSLP, PD-L1 and ENA78) were upregulated in all three conditions, whereas IL-12, MIP1α and IL-8 were specific for IL-1β stimulation, with IFNα2, IL-4, Il-15, IL-1β, MIP-1β, GIP and Ghrelin being selective for IL-1α stimulation, and IFNγ, MDC, IL-2, IL-5, IL-13, Eotaxin-3, TNFβ and IP-10 being selectively upregulated in response to LPS. Hence the PCKS platform can detect stimuli-selective as well as commonly induced inflammatory mediators.

**Figure 6.**
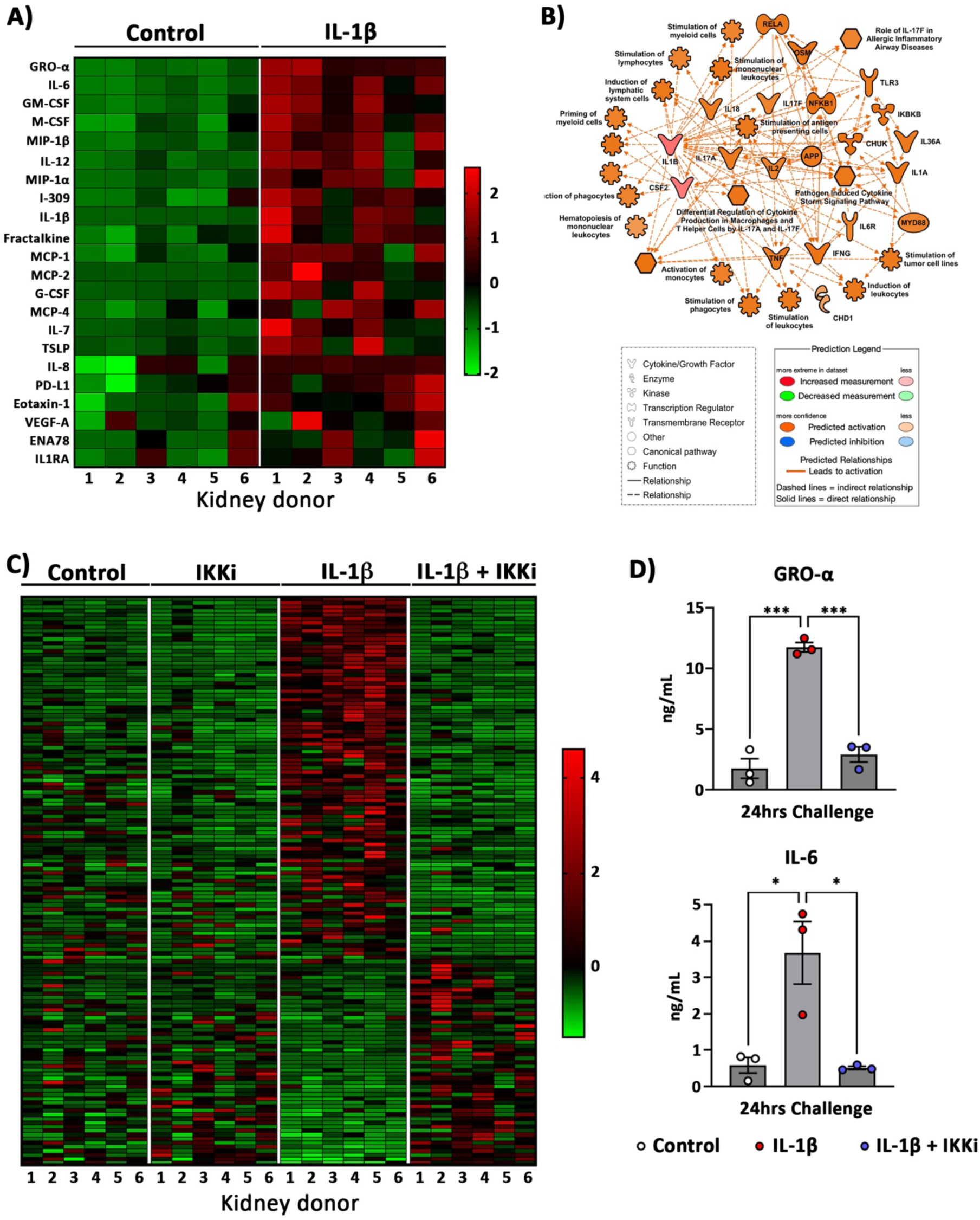
Human PCKS are responsive to inflammatory stimuli via activation of NF-κB. (A) Heat map of multiplex ELISA data showing secreted cytokines and chemokines in conditioned media from control and IL-1β-challenged PCKS (24 h challenge; n=6 donors). (B) Ingenuity Pathway Analysis interactome of proteins induced by IL-1β, identifying NF-κB pathway regulators and upstream activators. (C) Heat map of n=141 differentially expressed genes from bulk RNA-seq of PCKS after challenge with IL-1β in the presence or absence of an IKK2 inhibitor (IKKi) (n=5 donors). (D) Quantified secretion of GRO-α and IL-6 in PCKS media after a 24hrs IL-1β challenge ± IKKi. Data are mean ± SEM; *p<0.05, ***p<0.001.

IPA analysis revealed that these alterations constituted an interactome suggestive of the activation of multiple proinflammatory signalling pathways and identified NF-κB pathway regulators (e.g. NFKB1, IKBKB, RELA) and their upstream activators (e.g. TLRs, MYD88, TNF, IL6) as likely regulators of these processes (Figure 6B, Figure S9A-B and S9C-D). To test this hypothesis, we next treated PCKS with an IKKi (IKK2 inhibitor VI) to ask if blocking activation of NF-κB can suppress IL-1β-induced inflammation in PCKS. Bulk RNAseq on RNA isolated from PCKS revealed induction of 141 genes in response to challenge with IL-1β, with this induction attenuated by treatment with an IKKi (Figure 6C, Supplementary Table 7). Complementary KEGG, Reactome and Gene Ontology enrichment analyses of the IL-1β-induced genes confirmed activation of a coordinated NF-κB-driven inflammatory programme of cytokine and chemokine signalling, including IL-17 and Toll-like receptor pathways, and leukocyte chemotaxis (Supplementary Figure S10). Similarly, IL-1β induced secretion of GRO-α and IL-6 was also attenuated by IKKi (Figure 6D). These data highlight the potential for PCKS to be used for anti-inflammatory drug discovery in renal disease.

### Modelling fatty kidney disease in human PCKS reveals lipid-induced inflammation

We were next interested to ask the broader potential for PCKS in renal disease modelling. Lipid-mediated nephrotoxicity, part of which may reflect direct toxicity of high concentrations of circulating lipids in addition to secondary haemodynamic effects, has long been recognised as a cause of glomerular injury, however with the recent dramatic increase in incidence of obesity, the condition known as fatty kidney disease (or obesity-related glomerulopathy) is becoming a major cause of chronic kidney disease.^12^ In addition to the accumulation of renal ectopic fat, lipids can also accumulate within the renal parenchyma which is more likely to lead to tubular injury and secondary glomerulosclerosis.^13,14^ Given the increasing importance of fatty kidney disease as a cause of CKD we were interested to determine if PCKS can provide a model for this condition.

PCKS were cultured in the presence of increasing mM concentrations of a cocktail of lipids relevant to human metabolic disease^15,16^. As shown in Figure 7A, fatty acid treatment resulted in accumulation of fatty deposits within the renal parenchyma, with deposits appearing predominantly localised to tubular cells, at 2 mM, as demonstrated by Oil Red O staining. Targeted lipidomics showed a dose-dependent shift in lipid profile towards increased triacylglycerols (TAGs) and diacylglycerols (DAGs), indicative of fat storage in lipid droplets, alongside a progressive decline in phosphatidylcholine (PC) levels, consistent with shunting of lipid synthesis towards TAGs and/or stress-induced PC catabolism (Figure 7B, Supplementary Table 8). Consistent with this, principal component analysis (PCA) of the global lipidome showed clear separation of lipid-challenged PCKS from control and T0 samples, driven principally by a family of triacylglycerol species (including TG(52:1), TG(54:3), TG(54:4), TG(52:2) and TG(56:5)) together with PC(34:1), PC(36:4) and PE(38:4) (Figure 7C). These observations are indicative of incipient steatosis which can cause epithelial cell injury, oxidative stress and inflammation. Whilst DAGs promote activation of PKC, in diabetic kidney disease DAG-dependent aberrant activation of PKC has been linked to vascular complications, increased oxidative stress and renal inflammation.^17,18^ Moreover, a decline in PC levels is predictive of risk of developing end-stage kidney disease and poorer outcomes.^19,20^

**Figure 7.**
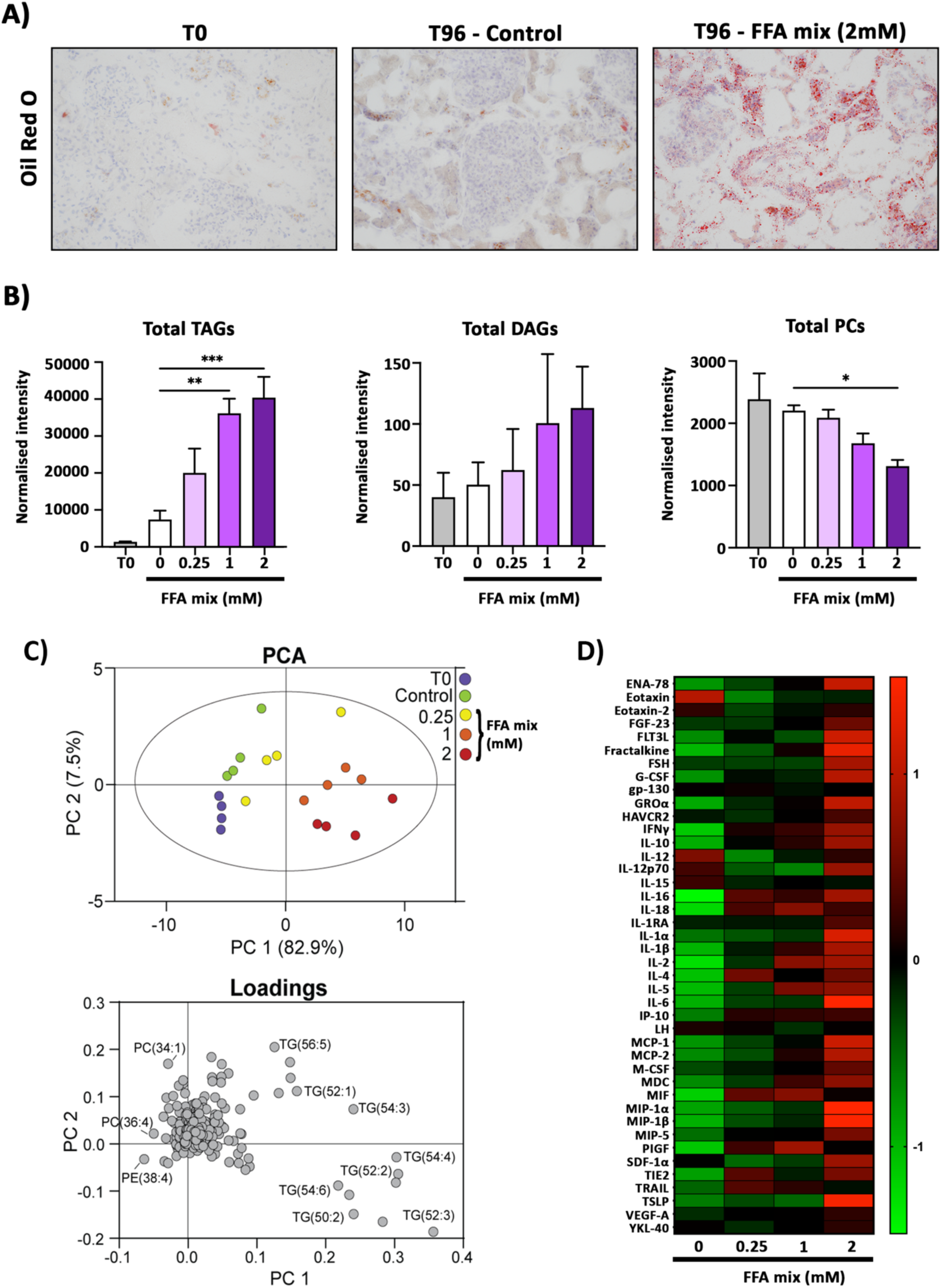
Modelling fatty kidney disease in human PCKS. (A) Representative Oil Red O staining of PCKS at T0, T96 control and T96 after challenge with 2 mM lipid, showing accumulation of neutral lipid within the renal parenchyma. (B) Mass spectrometry-based lipidomics (Supplementary Table 8) measured total triacylglycerols (TAGs), diacylglycerols (DAGs) and phosphatidylcholines (PCs) across increasing lipid concentrations (0, 0.25, 1, 2 mM); data are mean ± SEM (n=4 donors); *p<0.05, **p<0.01, ***p<0.001. (C) PCA scores plot of lipidomic profiles of fresh (T0), 96hrs control and lipid-challenged PCKS, with corresponding loadings plot identifying triacylglycerol species (including TG(52:1), TG(54:3), TG(54:4), TG(52:2) and TG(56:5)) together with PC(34:1), PC(36:4) and PE(38:4) as the principal contributors to sample separation. (D) Heat map of 43 cytokines, chemokines and growth factors quantified by multiplex ELISA (MSD) across increasing lipid concentrations.

Media lactate dehydrogenase (LDH) levels remained comparable between control and fatty acid-treated PCKS, indicating that lipid challenge did not cause overt cytotoxicity (Figure S11A).

Progression of steatosis to inflammation is a risk for perpetuation of renal damage and establishment of obesity-related CKD which has been reported to have caused in excess of 480 thousand deaths and 10 million DALYs globally in 2021.^21^ As inflammatory activation can occur independently of frank cell death, we used the MSD multiplex assay to measure secretion of inflammatory proteins into the PCKS media. Fatty acid treatment for 96hrs exposure was associated with dose-dependent increases in the expression of 43 different cytokines, chemokines and growth factors related to inflammatory processes (Figure 7D, Figure S11B). From these data we conclude that, while fatty acid treatment of human PCKS did not cause overt cytotoxicity, it did induce an inflammatory response, indicating that human PCKS can serve as a model system to study lipid-induced renal inflammation that underpins the development of obesity-related metabolic kidney disease. This highlights the potential of a fatty PCKS model to explore disease mechanisms and examine the therapeutic activities of metabolic modulators and anti-inflammatory drugs.

## Discussion

A significant challenge for biotechnology and pharmaceutical companies developing novel drugs for the treatment of kidney disease is to select pre-clinical testbeds that provide a translational bridge between the laboratory and the patient. Animal models, usually small rodent, are the mainstay for this purpose but are limited by inherent species differences that reduce their predictive value, plus there is increasing ethical considerations of the use of animals for drug discovery. 2D *in vitro* cell culture models are convenient and can be physiologically enhanced by generating specific cell types from human stem cells.^22^ However, 2D cell cultures fail to recapitulate the complex multi-cellular 3D tissue microenvironments and functions of the human kidney.^23^ Bioprinted kidney-on-a-chip systems and self-organising kidney organoids have received considerable attention, the latter arguably being the most advanced *in vitro* system for disease modelling and drug screening.^22^ However, the field still lacks an *in vitro* model that accurately recapitulates the biology of the human kidney and that takes into account numerous variables relevant to predicting drug efficacies and toxicities which include gender, the impacts of ageing, genetic factors and lifetime exposures of the kidney to dietary, metabolic and environmental factors.

Human PCKS generated from unused donor kidneys offer a viable *ex-vivo* alternative to bioprinted or organoid-derived 3D models as pre-clinical testbeds based on their faithful representation of the complexity of renal architecture.^3,4,22,24,25^ Moreover, with availability of anonymised donor information such as sex, age, BMI, smoking, alcohol use etc, PCKS can account for a wide variety of physiological and pathophysiological variables that impact on renal function and response to drugs. A variety of culture systems have been described for the maintenance of precision cut tissue slices,^26^ with the goal of ensuring cell viability and maintenance of tissue architecture and metabolic function over extended periods of culture. Here we have shown that our patented rocked bioreactor system supports human PCKS viability for 5 or more days in continuous culture; direct comparison with static culture conditions was not performed in this study and will be required to establish the comparative benefit of the bioreactor system. This time window is sufficient to model spontaneous fibrosis, induced inflammation and fibrosis by relevant physiological (e.g. endotoxin, inflammatory mediators and lipids) triggers, and drug-induced acute damage. We have shown that PCKS offer the opportunity for spatial resolution of collagen fibre deposition into the renal parenchyma which can help unlock new anti-fibrotics that target the synthesis and/or maturation of collagen in the damaged kidney. Although not tested here, PCKS should also be well suited to spatial mapping of gene and protein expression, and we have demonstrated that PCKS can detect changes at transcriptome, proteome (cytokine/chemokine) and lipidome scale in PCKS responding to relevant stimuli such as fatty acid treatment and exposure to pro-fibrogenic growth factors. Further applications include examining the efficacy and associated toxicities of biologics, polynucleotide-based drugs (e.g. siRNA) and cell-based therapies.

While human PCKS offer many advantages listed above including meeting key 3Rs demands by potentially replacing the need for animal models, they do have several limiting factors. Sourcing donor kidneys requires considerable logistical considerations that can be a challenge for many academic as well as industrial laboratories. Maintaining tissue viability involves detailed characterisation of culture conditions to ensure optimal supply of oxygenation and nutrients to tissue slices features that our rocked bioreactor system is designed to address, including facilitating the removal of toxic metabolites, although this specific benefit has not been directly demonstrated in the present study. At the physiological level the lack of perfusion and a blood supply is a significant limitation that disallows systemic physiological factors and the influences of recruited immune cells to be studied. However, we have been able to demonstrate that local inflammatory signalling events involving tissue activation of NF-κB and the elaboration of complex cytokine and chemokine pathways can be recapitulated in PCKS. In addition, deceased-donor kidneys are exposed to warm and/or cold ischaemia before culture, which likely contributes to the spontaneous fibrosis observed but also offers an opportunity to test early interventions in renal injury and inflammation.

We conclude that human PCKS are an important physiological platform for exploring the biology of the human kidney and are a sufficient excerpt of the organ to lend themselves to the drug discovery process.

## Data Sharing Statement

The data supporting the findings of this study are available from the corresponding author upon reasonable request, subject to applicable ethical approvals, donor confidentiality requirements and relevant data-sharing agreements.

## Declaration of generative AI and AI-assisted technologies in the manuscript preparation process

During the preparation of this manuscript, the authors used OpenAI ChatGPT solely for language editing to improve grammatical accuracy, clarity, readability and consistency of expression. The tool was not used to generate scientific content, analyse data, develop interpretations or draw conclusions. The authors reviewed and edited the output as needed and take full responsibility for the content of the published article.

## Supplementary Figure Legends

**Figure S1.**
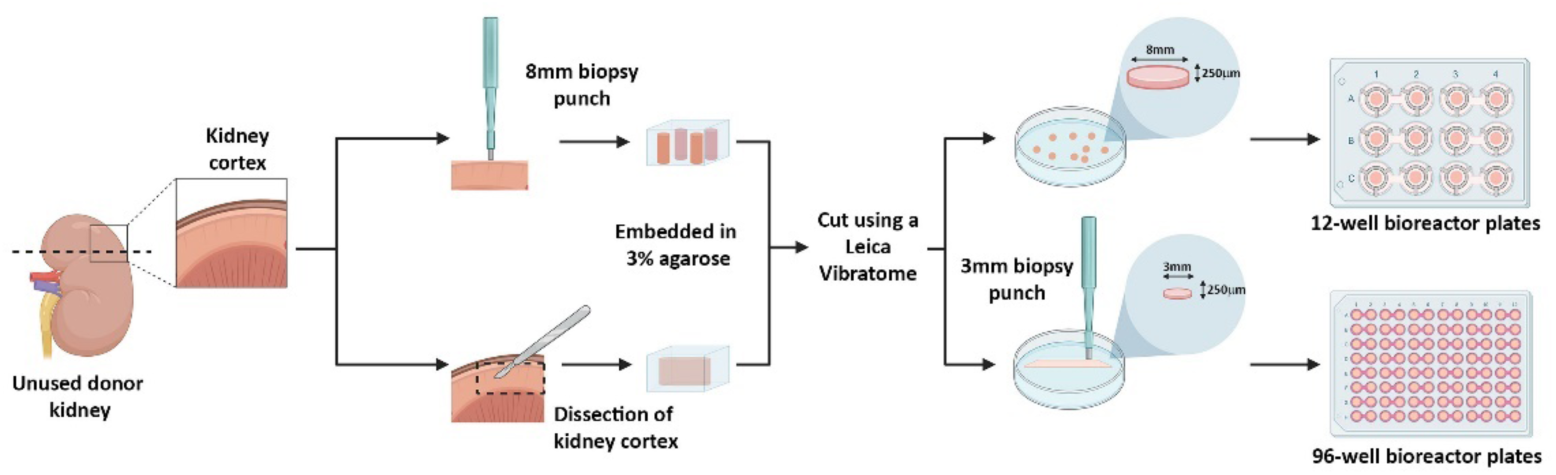
Generation of PCKS from human kidney. Sections of renal cortex are dissected from the kidney and cored with an 8 mm biopsy punch. The tissue is embedded in 3% agarose and sliced using a Leica VT1200S vibratome to produce 8 mm-wide, 250 µm-thick slices, which are placed into 8-µm Transwell inserts and cultured in a 12-well bioreactor plate. Alternatively, renal cortex is embedded directly in agarose to produce 250 µm-thick tissue sheets, cored with a 3 mm biopsy punch, placed into 8-µm Transwell inserts and cultured in a 96-well bioreactor plate.

**Figure S2.**
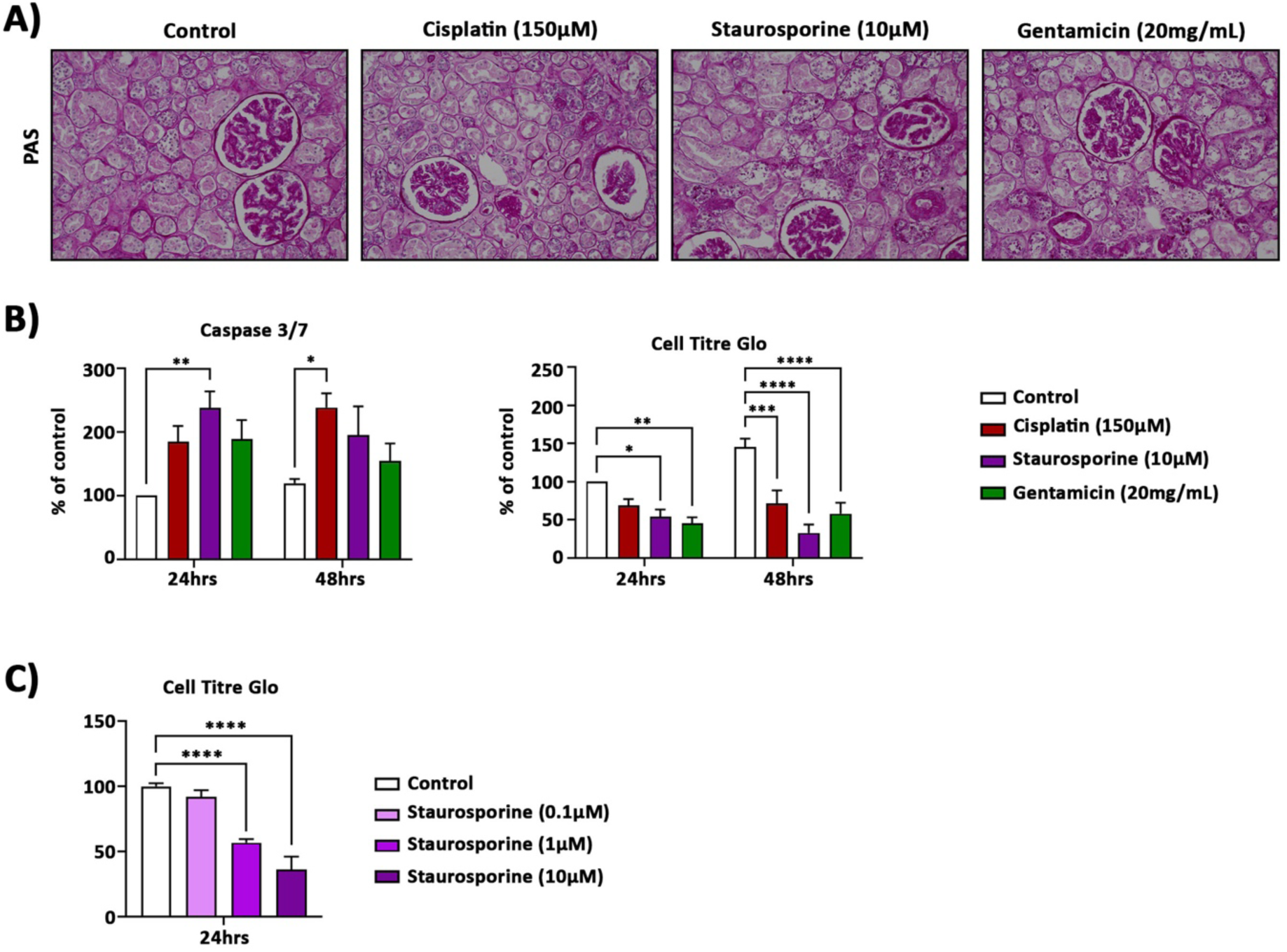
Modelling drug-induced acute renal damage in human PCKS. (A) Representative Periodic Acid-Schiff (PAS) staining of PCKS after 24hrs exposure to control, cisplatin (150 µM), staurosporine (10 µM) or gentamicin (20 mg/mL), showing drug-induced tubular injury. (B) Caspase 3/7 activity and Cell Titre-Glo in PCKS (n=4 donors) readout (% of control) at 24 and 48hrs across treatment groups. (C) Cell Titre-Glo dose–response to staurosporine (0.1, 1, 10 µM) at 24hrs. Data are mean ± SEM; *p<0.05, **p<0.01, ***p<0.001, ****p<0.0001.

**Figure S3.**
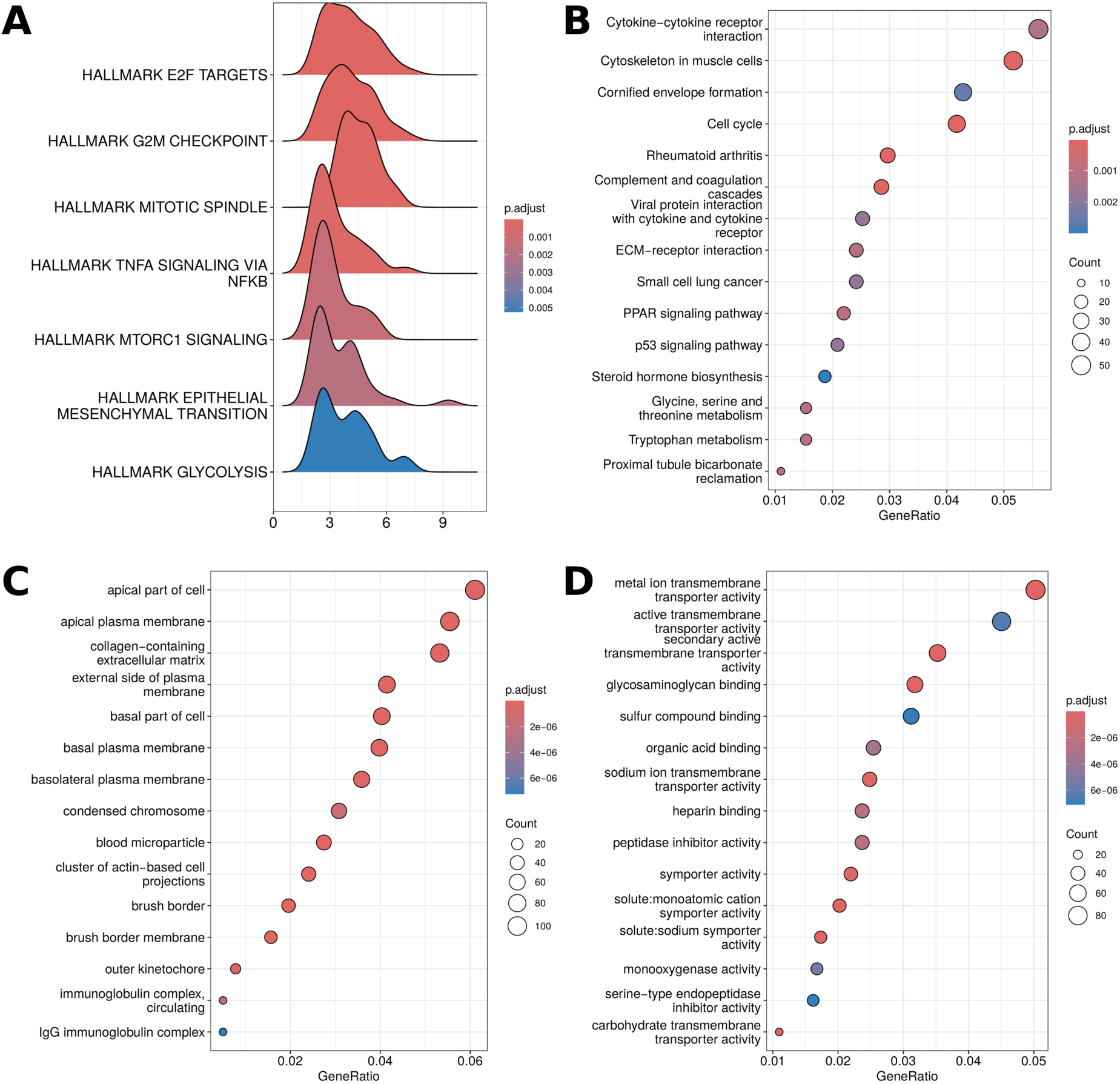
Complementary pathway enrichment analysis of genes differentially expressed during spontaneous fibrogenesis in human PCKS. Differentially expressed genes between T0 and T96 PCKS (n=5 donors; Figure 1C) were subjected to complementary enrichment analyses. (A) Gene set enrichment analysis (GSEA) against the MSigDB Hallmark collection, shown as a ridgeplot, with enriched programmes including epithelial–mesenchymal transition, TNFα signalling via NF-κB, E2F targets, G2M checkpoint, mitotic spindle, mTORC1 signalling and glycolysis. (B) KEGG pathway over-representation analysis, shown as a dotplot, spanning extracellular matrix and metabolic remodelling, inflammation, proliferation and kidney-specific transport and metabolic processes (proximal tubule bicarbonate reclamation, steroid hormone biosynthesis, amino-acid metabolism). (C) Gene Ontology Cellular Component (GO:CC) analysis highlighting tubular epithelial structure (apical, basal and basolateral plasma membrane, brush border) and collagen-containing extracellular matrix. (D) Gene Ontology Molecular Function (GO:MF) analysis highlighting renal transmembrane transport activities alongside matrix-binding and peptidase-inhibitor activities. For all dotplots, point size denotes gene count (Count) and colour the Benjamini–Hochberg adjusted p-value (p.adjust); x-axis shows GeneRatio.

**Figure S4.**
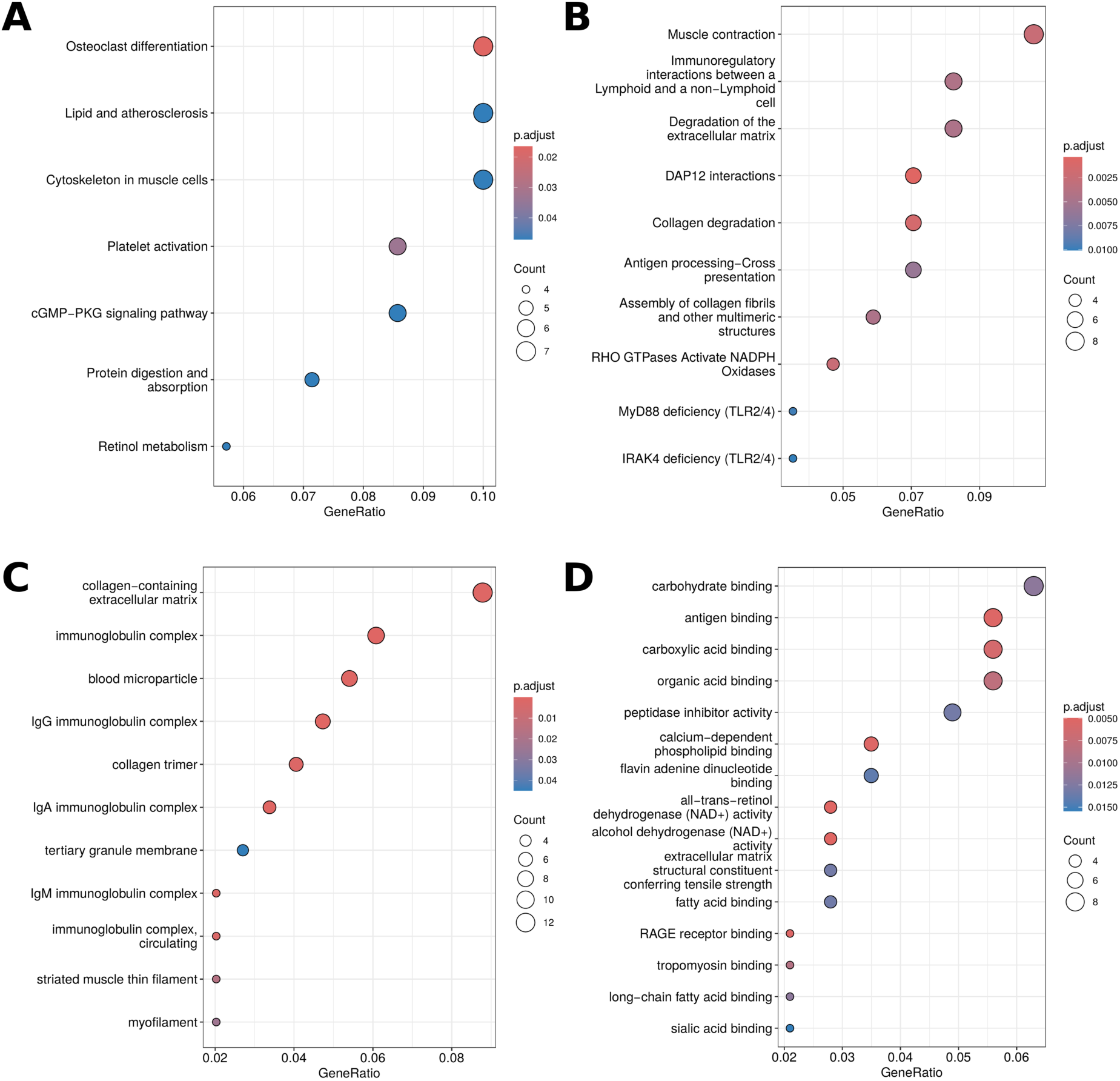
Complementary pathway enrichment analysis of genes modulated by anti-fibrotic treatment in human PCKS. Differentially expressed genes identified in the control vs. nintedanib comparison at T96 (n=5 donors; Figure 2E) were subjected to complementary over-representation analyses; pirfenidone was excluded from this analysis as it did not produce a comparable transcriptional response (see main text). (A) KEGG pathway analysis highlighting metabolic and lipid-handling pathways, cytoskeletal organisation and signalling/activation pathways. (B) Reactome analysis dominated by collagen and extracellular matrix remodelling (collagen degradation, degradation of the extracellular matrix, assembly of collagen fibrils) alongside innate immune and inflammatory signalling (DAP12 interactions, MyD88 and IRAK4 deficiency [TLR2/4]). (C) Gene Ontology Cellular Component (GO:CC) analysis highlighting collagen and matrix structures, immunoglobulin complexes and contractile structures. (D) Gene Ontology Molecular Function (GO:MF) analysis highlighting extracellular matrix structural constituents, lipid- and retinoid-related activities and RAGE/peptidase-inhibitor activities. Point size denotes gene count (Count) and colour the adjusted p-value (p.adjust); x-axis shows GeneRatio.

**Figure S5.**
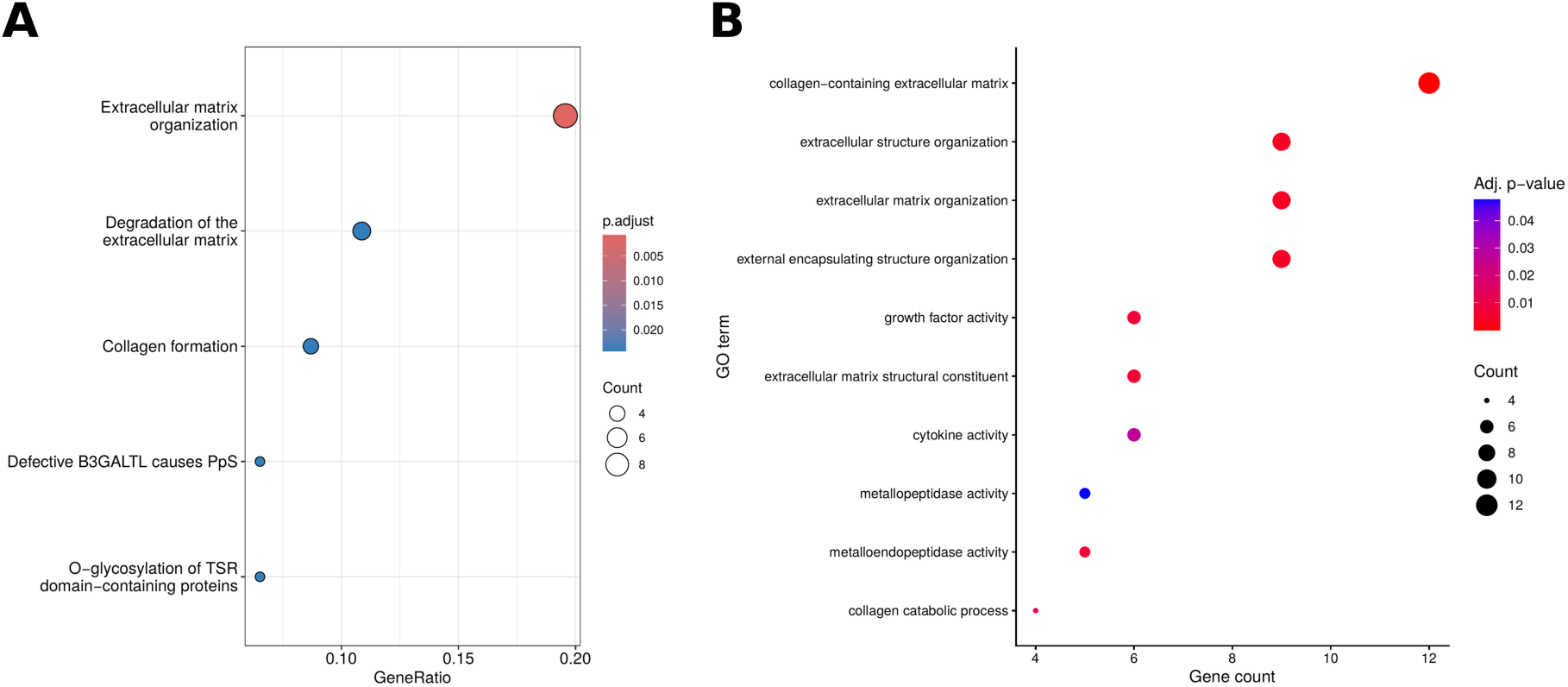
Complementary pathway enrichment analysis of TGFβ1-regulated genes in human PCKS. Differentially expressed genes between control, ALK5i, TGFβ1 and TGFβ1 + ALK5i groups at T96 (n=5 donors; Figure 3E) were subjected to complementary over-representation analyses. (A) Reactome pathway analysis demonstrating enrichment of extracellular matrix organisation, degradation of the extracellular matrix, collagen formation and collagen O-glycosylation. (B) Combined Gene Ontology analysis (Biological Process, Cellular Component and Molecular Function) highlighting collagen-containing extracellular matrix, extracellular matrix and structure organisation, collagen catabolic process, extracellular matrix structural constituent, metalloendopeptidase activity, growth factor activity and cytokine activity. Point size denotes gene count (Count) and colour the adjusted p-value (p.adjust).

**Figure S6.**
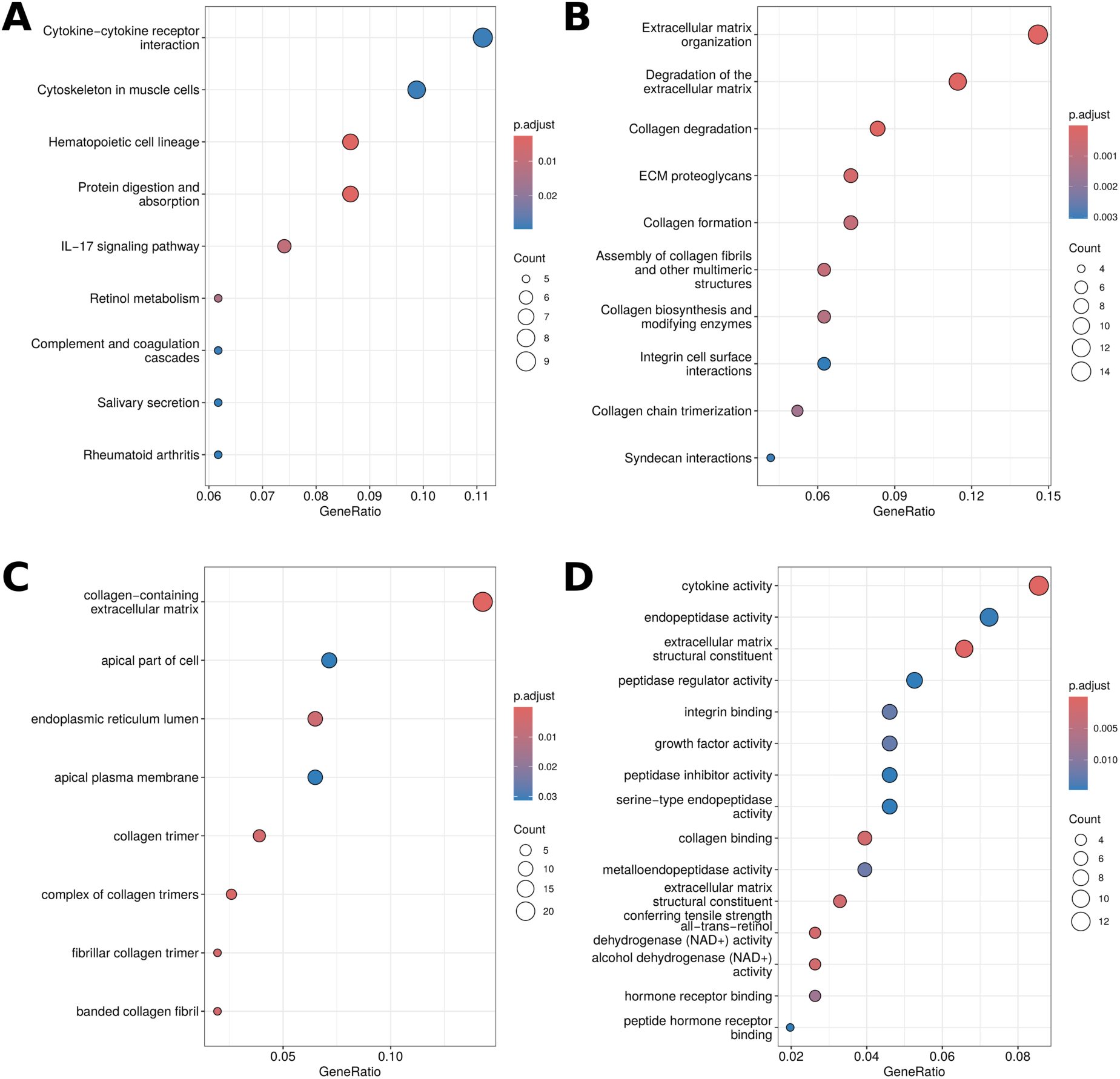
Complementary pathway enrichment analysis of genes modulated by anti-fibrotic treatment of TGF-β1-stimulated human PCKS. Differentially expressed genes identified in the TGF-β1 vs. TGF-β1 + nintedanib comparison at T96 (n=5 donors; Figure 5E) were subjected to complementary over-representation analyses; TGF-β1 + pirfenidone was excluded from this analysis as it did not produce a comparable transcriptional response (see main text). (A) KEGG pathway analysis highlighting inflammatory and immune pathways, metabolic pathways and cytoskeletal organisation. (B) Reactome analysis dominated by collagen and extracellular matrix remodelling alongside cell–matrix adhesion (integrin and syndecan interactions). (C) Gene Ontology Cellular Component (GO:CC) analysis highlighting collagen and matrix structures and secretory/epithelial compartments. (D) Gene Ontology Molecular Function (GO:MF) analysis highlighting cytokine and growth factor activity, extracellular matrix structural constituents, collagen and integrin binding and a peptidase programme. Point size denotes gene count (Count) and colour the adjusted p-value (p.adjust); x-axis shows GeneRatio.

**Figure S7.**
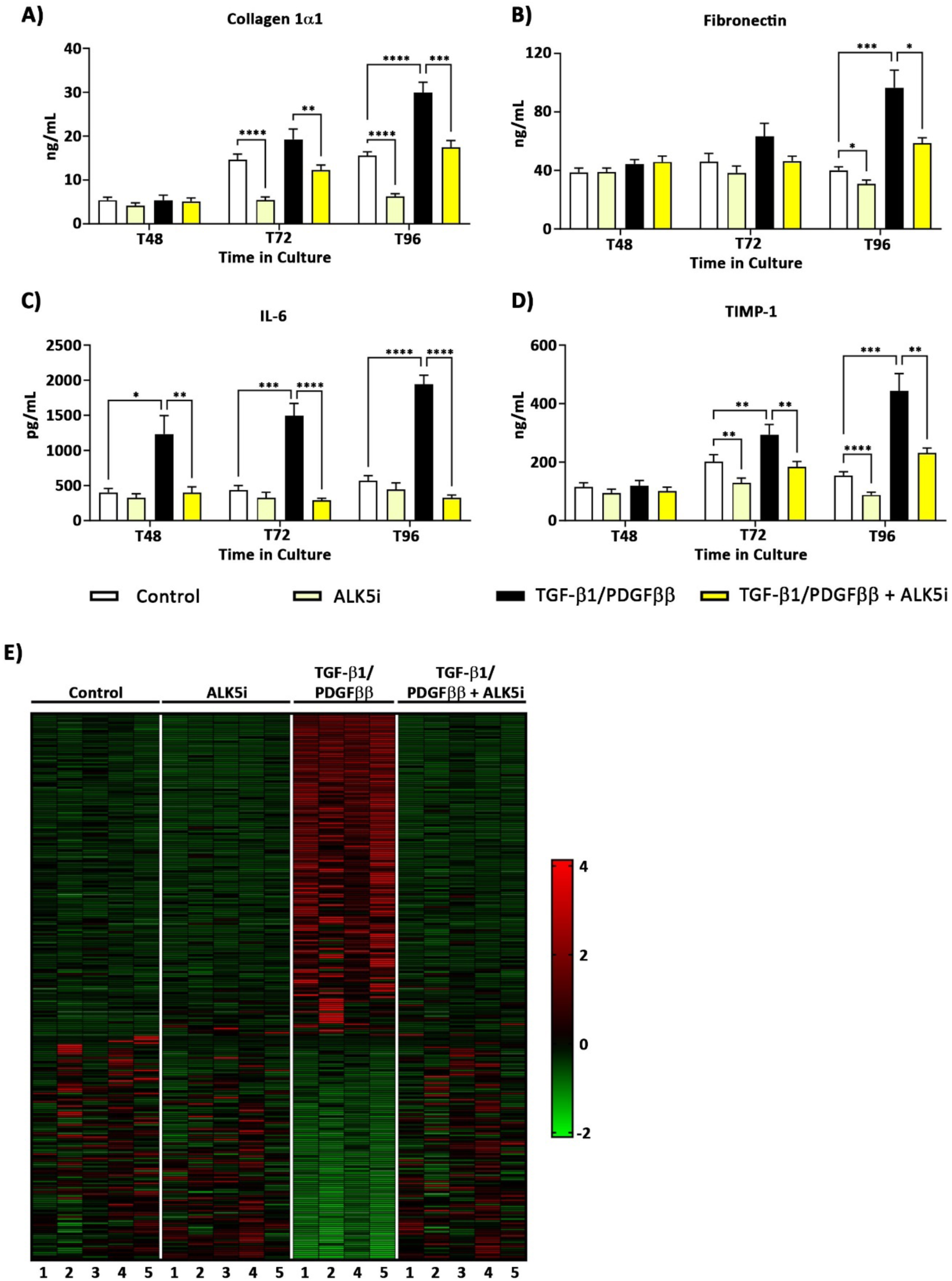
Inducing fibrosis with combined pro-fibrogenic stimuli TGF-β1/PDGFββ in human PCKS. (A–D) Quantified levels of collagen 1α1, fibronectin, IL-6 and TIMP-1 in PCKS culture media after 48hrs (T48), 72hrs (T72) and 96hrs (T96) in the control group and in the presence of ALK5i, TGF-β1/PDGFββ, or TGF-β1/PDGFββ + ALK5i. Data are mean ± SEM (n=6 donors), analysed by two-way ANOVA with Dunnett’s multiple comparisons; ****p<0.0001. (E) Heat map of n=334 differentially expressed genes between control, ALK5i, TGFβ1/PDGFββ and TGFβ1/PDGFββ + ALK5i groups at T96 (n=5 donors).

**Figure S8.**
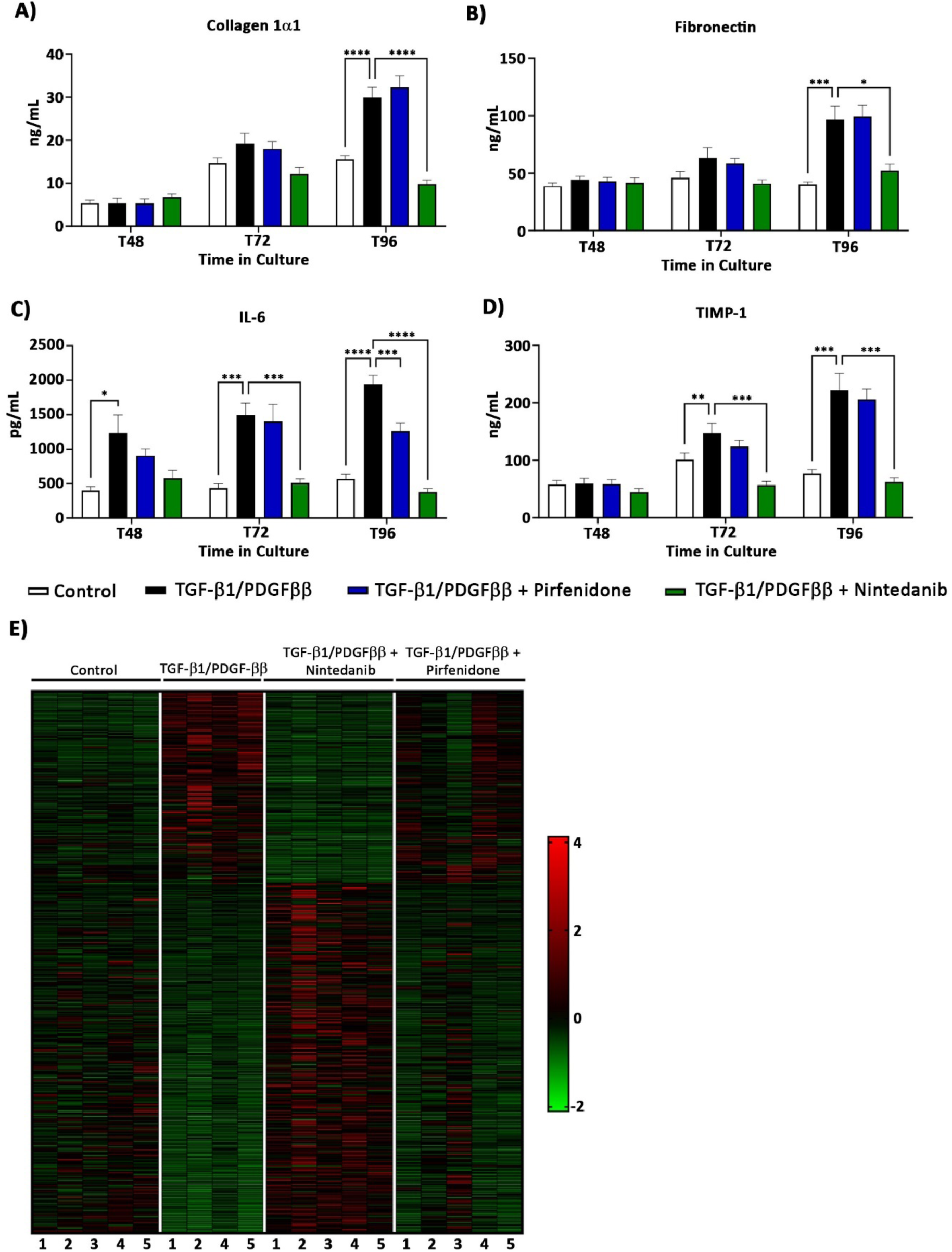
Testing clinically approved anti-fibrotics in human PCKS in combination with pro-fibrotic TGF-β1/PDGFββ stimuli. (A–D) Quantified levels of collagen 1α1, fibronectin, IL-6 and TIMP-1 in PCKS culture media after 48hrs (T48), 72hrs (T72) and 96hrs (T96) in the control group and in the presence of TGF-β1/PDGFββ, nintedanib or pirfenidone. Data are mean ± SEM (n=6 donors), analysed by two-way ANOVA with Dunnett’s multiple comparisons; ****p<0.0001. (E) Heat map of differentially expressed genes between control, TGFβ1/PDGFββ, nintedanib and pirfenidone groups at T96 (n=5 donors).

**Figure S9.**
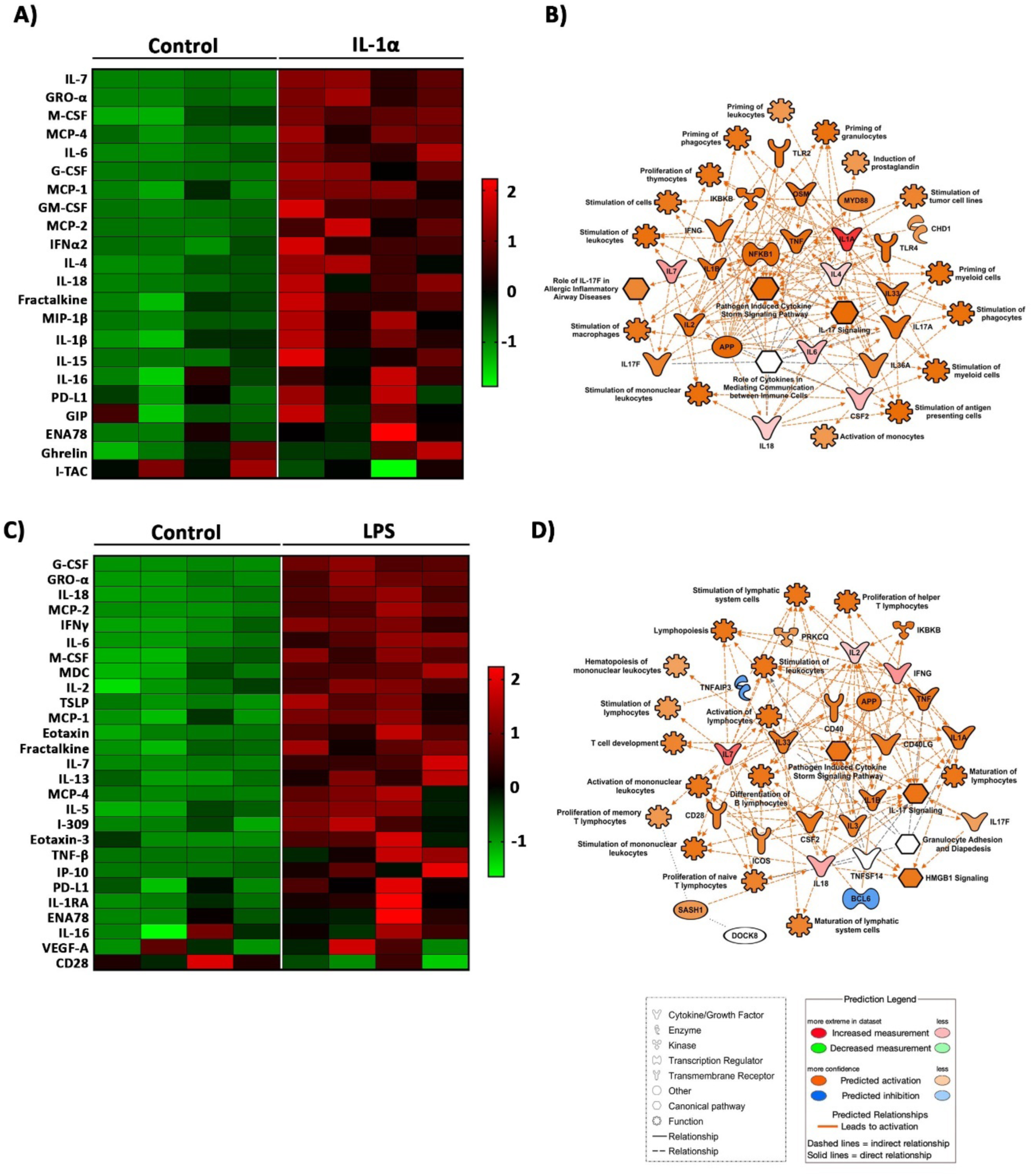
Human PCKS responses to IL-1α and LPS. PCKS were challenged for 24hrs with IL-1α or LPS and secreted cytokines and chemokines in the conditioned media were quantified by multiplex ELISA. (A) Heat map of secreted cytokines and chemokines in conditioned media from control and IL-1α-challenged PCKS (n=6 donors). (B) Ingenuity Pathway Analysis interactome of proteins induced by IL-1α, identifying NF-κB pathway regulators and upstream activators. (C) Heat map of secreted cytokines and chemokines in conditioned media from control and LPS-challenged PCKS (n=6 donors). (D) Ingenuity Pathway Analysis interactome of proteins induced by LPS, identifying NF-κB pathway regulators and upstream activators.

**Figure S10.**
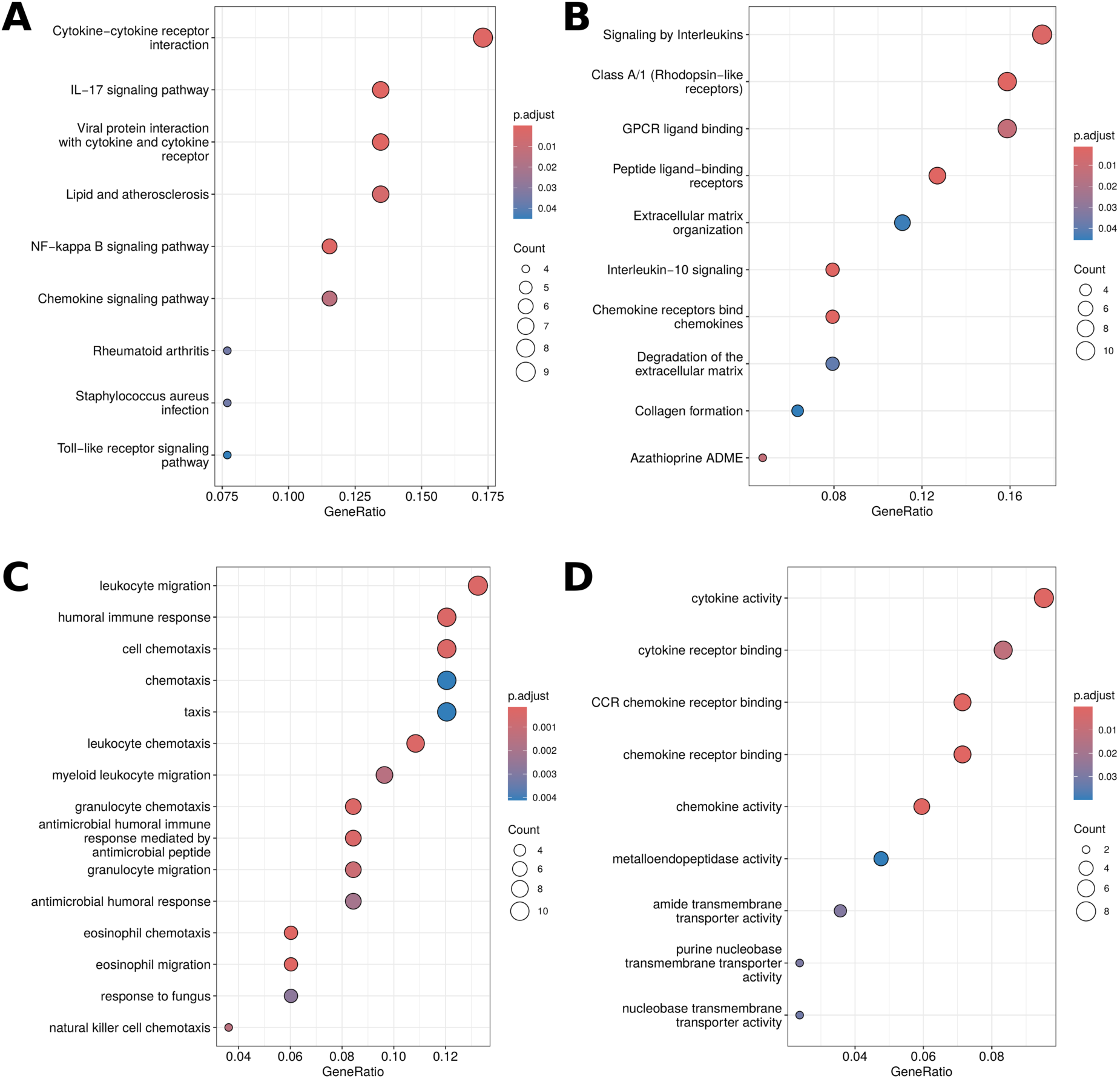
Complementary pathway enrichment analysis of IL-1β-induced, NF-κB-dependent genes in human PCKS. Differentially expressed genes induced by IL-1β challenge and attenuated by an IKK2 inhibitor (IKKi) at T96 (n=5 donors; Figure 6C) were subjected to complementary over-representation analyses. (A) KEGG pathway analysis demonstrating enrichment of core inflammatory signalling pathways including NF-κB signalling, IL-17 signalling, Toll-like receptor signalling, cytokine–cytokine receptor interaction and chemokine signalling. (B) Reactome analysis highlighting interleukin signalling and chemokine and peptide-ligand receptor signalling. (C) Gene Ontology Biological Process (GO:BP) analysis dominated by leukocyte recruitment and chemotaxis and humoral and antimicrobial immune responses. (D) Gene Ontology Molecular Function (GO:MF) analysis highlighting cytokine and chemokine activity and receptor binding alongside metalloendopeptidase activity. Point size denotes gene count (Count) and colour the adjusted p-value (p.adjust); x-axis shows GeneRatio.

**Figure S11.**
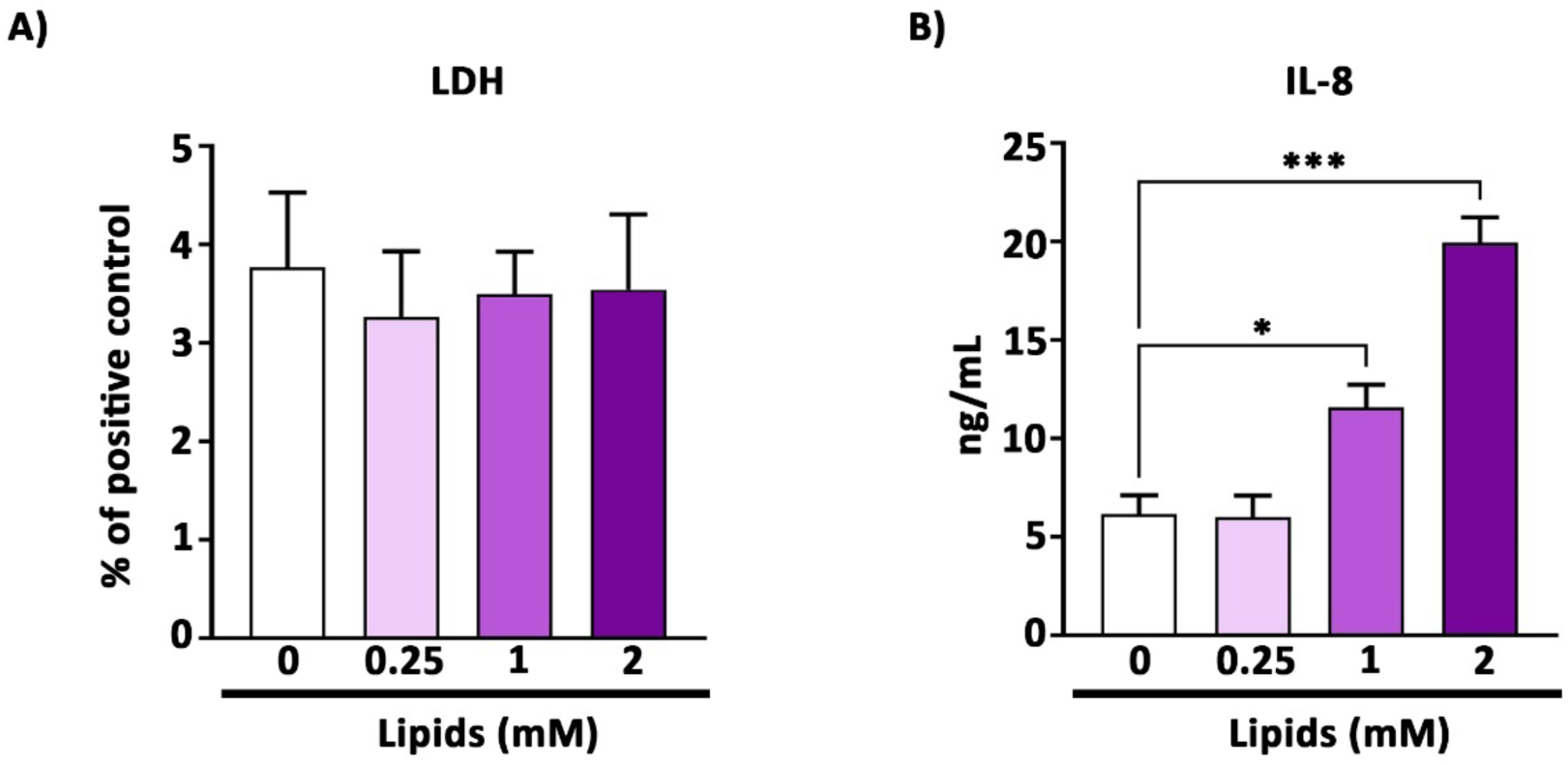
Lipid challenge does not cause overt nephrotoxicity and induces IL-8 secretion. (A) Media lactate dehydrogenase (LDH) levels in control and fatty acid-treated PCKS (n=4 donors) across increasing lipid concentrations (0, 0.25, 1, 2 mM). (B) IL-8 in PCKS media confirming dose-dependent increases at 96 h of lipid exposure (0, 0.25, 1, 2 mM); *p<0.05, ***p<0.001.

