## Supplementary materials and methods for "Human Precision-Cut Kidney Slices as a Translational Platform for Modelling Fibrotic, Inflammatory, Metabolic and Drug-Induced Kidney Injury"

**Ethical Approval**

Human kidneys from deceased UK donors were obtained through the Newcastle Institute of Transplantation Tissue Biobank. Consent for organ donation and research was obtained by Specialist Nurses in Organ Donation in accordance with national organ donation procedures. The Newcastle Institute of Transplantation Tissue Biobank operates under approval from an NHS Research Ethics Committee (REC reference: 22/NE/0064) for the collection, storage and distribution of human tissue for research, with delegated authority to release tissue to approved research studies. All samples used in this study were provided through this governance framework.

**Preparation and ex-vivo culture of Precision Cut Kidney Slices**

*12 well* - Tissue cores from the cortex of the human kidney were prepared using 8mm Stiefel biopsy punch (Medisave, Weymouth, UK), and PCKS were generated using a Leica VT1200S vibrating blade microtome, then cultured in 8µm pore Transwell inserts (665638, Greiner), in modified tissue culture plates and rocked on a bioreactor platform as previously described.[1,2] PCKS were cut in DMEM/F12 (1:1) - (21331-020, Gibco) supplemented with 1% (v/v) penicillin/streptomycin (P4333 - Sigma-Aldrich), 1% (v/v) L-glutamine (G7513 - Sigma-Aldrich), 1% (v/v) Sodium pyruvate (S8636, Sigma-Aldrich) and 1x REGM™ Singlequots™ Supplement Pack (CC-4127, Lonza), and then cultured in the same media in the bioreactor platform at 37°C, supplemented with 21% $O_2$ and 5% $CO_2$. Media was changed daily for the duration of culture.

*96 well* - Kidney cortex was dissected into ~1cm$^3$ tissue blocks, transferred to a metal mould, submerged in 3% low gelling temperature agarose (A9414; Sigma-Aldrich, Poole, UK), and placed on ice for 5 minutes. The agarose block was cut as previously described[1] to generate 250µm thick sheets of tissue and a 3mm Stiefel biopsy punch (Medisave, Weymouth, UK) was used to generate PCKS. PCKS were placed in 8µm pore, 96-well transwell inserts (3384, Corning, Corning) and cultured as described above (Supplementary Figure S1).

**Modelling inflammation and fibrosis in PCKS**

*Fibrosis* - PCKS were rested for 24hrs and then stimulated with TGF-β1 (10ng/mL,100-21, Peprotech) alone or in combination with PDGF-ββ (50ng/mL, 100-14B, Peprotech) for a further 72hrs to induce fibrosis. PCKS were stimulated in the presence or absence of activin receptor-like kinase 5 inhibitor (ALK5i) (SB-525334, S8822, Sigma-Aldrich), Nintedanib (2.5μM, 11022, Cayman Chemical) or Pirfenidone (2.5mM, 13986, Cayman Chemical). Stimuli and inhibitors were refreshed daily for the duration of the experiment. PCKS were harvested for immunohistochemistry or downstream RNA isolation & sequencing as described below.

*Inflammation* - PCKS were rested for 24hrs and then given a single challenge with IL-1α (1ng/mL, 200-01B, Peprotech), IL-1β (1ng/mL, 200-01B, Peprotech) or LPS (1μg/mL, L4391, Sigma-Aldrich) for a further 24hrs to induce inflammation. PCKS were stimulated in the presence or absence of an IKK2 inhibitor (IKK2 Inhibitor VI) (10μM, 17276, Cayman Chemical).

**PCKS lipid treatment**

After a 24hr rest period, PCKS were challenged with were treated with 0.25 mM, 1 mM or 2 mM free fatty acid mixture in 2.6% fatty acid-free bovine serum albumin (BSA) (A8806, Sigma-Aldrich) for 72hrs. Lipids were refreshed daily for the duration of the experiment.

**Toxicity induction**

After a 48hr rest period, PCKS were challenged with cisplatin (150μM, #2251, Tocris), staurosporine (10μM, #1285, Tocris) or Gentamicin Sulfate (20μg/ml, #G3632, Sigma-Aldrich) for 24hrs. Media and tissue samples were collected for soluble outputs, Glo assays or histology, at either directly after the challenge (24hrs) to assess peak injury, or PCKS were changed into fresh media and cultured for a further 24hrs to assess recovery from injury.

For all PCKS experiments media with or without stimuli and/or compounds were changed daily for the duration of the experiment and tissue culture supernatant was collected daily after the 24hrs rest period for the quantification of soluble markers.

**Immunohistochemistry**

Formalin fixed, paraffin embedded (FFPE) PCKS were cut at 5μm thickness and processed for haematoxylin and eosin (H&E), Picrosirius Red and αSMA staining as previously described.[3] Periodic Acid-Schiff (PAS) stain was performed on renal sections using a PAS staining kit (ab150680) according to the manufacturer's instructions. Representative images were acquired at x10 or x20 magnification using a Nikon ECLIPSE Ni-U microscope.

**Histology**

FFPE tissue sections were stained with Picrosirius Red to assess collagen deposition. Whole-slide digital images were acquired at ×40 magnification (0.23 μm/pixel) using a AxioScan 7 (Zeiss, UK) whole slide imager.

**Digital Pathology Analysis**

FibroNest™ (PharmaNest Inc, Princeton, NJ, USA)[4,5], a cloud based, high-resolution, single-fiber image analysis platform was used to quantify the fibrosis phenotype in the context of three complementary phenotypic layers for (i) collagen deposition and structure features (12 traits), (ii) fiber morphometry (12 traits), and (iii) fibrosis architecture (7 traits to measures the organization of the fibers[6,7]). Color normalization and standardization[8,9] was performed to calibrate the images of the study. Each trait was quantified with 7 quantitative parameters (qFTs) to account for severity, distortion, and variance, resulting in a total of 313 qFTs, one of them being Collagen Area Ratio. Fibers were also classified into "fine" (simple skeleton or low node/branch ratio) and "assembled" (complex skeleton or high node/branch ratio). The qFT dataset was automatically (AI) surveyed to identify traits (principal qFTs) that would exhibit a significant ($p<0.05$) and meaningful (>20%) relative difference (group average) between the control ("Control" group) and the group that most expresses the disease phenotype ("TGF-b1" group). Such principal qFT and their variation can be visualized in the form of a heat chart[10] and assembled into an equi-weighted and normalized Phenotypic Fibrosis Composite Score (Ph-FCS)[11]. Augmented digital pathology images help visualize any of these measurements.

Similarly, the principal qFT that are associated with a specific phenotypic layer can be assembled in a specific score, for instance the Fibrosis Morphometric Composite Score for Fine fibers. Here, we analyzed the full tissue area of the tissue section.

A workflow of the method, the full list of qFTs and the analytical performance of the method has been published elsewhere[4]. This automated method has superior detection thresholds to detect biologically relevant differences in the histological phenotype of fibrosis, due to disease etiology[4], severity across different biological models[12], treatment response[11,13] or outcomes prediction[14]. Most recently, the fusion of such single-fiber datasets with spatial single-cell RNA-seq data[15] demonstrated that the changes in the morphometric phenotypes of fibrosis captured by the FibroNest approach correspond to underlying biological remodeling - a key finding that bridges digital pathology with cell-level biology and substantiate the utility of the approach.

**ELISA**

Levels of human Collagen 1α1 (DY6220), TIMP-1 (DY970), Hyaluronic Acid (DY2089), Fibronectin (DY1918), IL-6 (DY206), MCP-1 (DY279), CXCL-1/GRO-α (DY275), IL-8/CXCL8 (DY208), or HSP70/HSPA1A (DY1663-05) in the tissue culture supernatant were quantified using Duoset ELISA kits (R&D Systems) as per manufacturer's instructions. A multi-plex ELISA (U-PLEX Immuno-Oncology Group 1 (hu) 111-Plex - K15342K-1, Meso Scale Discovery (MSD)) was used to quantify secreted proteins released from PCKS in response inflammatory stimuli as per manufacturer's instructions.

**Colorimetric & Luminescent Assays**

CyQUANT™ LDH Cytotoxicity Assay (C20300, Thermo Fisher), Cell Titer-Glo® 3D Cell Viability Assay (G9681, Promega), CellTox™ Green (G8741, Promega), Caspase 3/7-Glo® 3D Assay (G8981, Promega) were all performed according to the manufacturers' instructions. Resazurin assay was performed as previously described[16].

**RNA isolation and sequencing**

PCKS (n=2 PCKS per group) were suspended in QIAzol in 2mL tubes containing 2.8mm ceramic beads (432-3752, Precellys), homogenised using at Precellys24 tissue homogeniser (Bertin Technologies, Montigny-le-Bretonneux, France) and passed through a Qiashredder (79656, Qiagen). Next, chloroform was added, the sample vortexed and centrifuged at 12,000g for 15 minutes. The aqueous layer was collected, added to 70% ethanol and total RNA isolated using the RNeasy Micro Kit as per manufacturer's instructions (74004, Qiagen). Total RNA quality was assessed prior to library preparation. RNA-seq libraries were prepared at the Newcastle University Genomics Core Facility (Newcastle, UK) using a stranded mRNA library preparation protocol. Libraries were sequenced to generate 75 bp single-end reads, with a sequencing depth of >30 million reads per sample. Sequencing quality was assessed by FastQC (Available online at http://www.bioinformatics.babraham.ac.uk/projects/fastqc/). Sequences were then trimmed to eliminate poor-quality bases and adaptors using Trimmomatic software [17] allowing a maximum of 1 mismatch and an average quality threshold of 30. The *quasi-mapping* mode of Salmon[18] was used for the mapping and quantification of the transcripts. The DESeq2 R package[19] was used to normalize read counts by scaling a normalized factor in all libraries and to assess the differential expression. The *P* values were corrected by the Benjamini and Hochberg method[20]; then, the expression level was considered significantly different at *P* adjusted of <.001 and log2 fold change of >2. Plots were created with gplots R package. Differentially expressed genes as selected by R were input into the IPA tools (http://www.qiagenbioinformatics.com/products/ingenuity-pathwayanalysis/) for functional annotation.

**Oil Red O staining**

A 0.5% stock solution was made by dissolving Oil Red O (C.I. 26125, Sigma) in 100% isopropanol, then diluted in a 3:2 ratio with deionised water to create a 60% Oil Red O working solution. The solution was filtered prior to use. 10 µm sections of cryo-frozen PCKS tissue were generated and formalin fixed for 30 minutes. The sections were then washed with running deionised water for 2 minutes, then rinsed with 60% isopropanol for 2 minutes. The sections were then stained in freshly prepared Oil Red O working solution for 15 minutes, then rinsed with 60% isopropanol for 1.5 minutes, before being counterstained with Mayer's Haematoxylin. The sections were rinsed in deionised water and mounted with Glycergel (C0563, Dako).

**Bulk lipidomics - Lipid extraction**

PCKSs were homogenised in chloroform: methanol (2:1 v/v, 1 mL) using Precellys tissue homogeniser (Bertin Technologies, Montigny-le-Bretonneux, France). Lipid extraction was performed using the Folch method[21]. Briefly, LC-MS grade water (400 µL) was added to the homogenised PCKSs in organic solvent and thoroughly mixed. Separation of the polar and non-polar fractions was achieved by centrifugation (13,200 g, 15 min). The separate fractions were dried down under vacuum. The organic layer was reconstituted in 400 µL chloroform: methanol (2:1 v/v) and diluted (1:20 v/v) with 2:1:1 isopropanol: acetonitrile: water, with the addition of an internal standard mix composed of isotopically labelled compounds from major lipid classes (Avanti Polar Lipids, Inc., Alabaster, AL USA).

**Mass spectrometry**

Sample (5 µL) was injected onto a C18 BEH column (2.1 x 50 mm,1.7 µm; Waters Ltd, Wilmslow, U.K.) and separated using ultra-high performance liquid chromatography on a Accela 3000 UHPLC (Thermo Fisher Scientific) using gradient elution at 0.5 mL/min and column temperature 55 °C. Mobile phase A was 60:40 acetonitrile: water and mobile phase B was 90:10 isopropanol: acetonitrile, both containing 10 mM ammonium formate Starting mobile phase was 40% B, increasing to 99% B over 8.5 minutes, before returning to starting conditions for 1.5 minutes. UHPLC was coupled to a LTQ Orbitrap Eilte (Thermo Fisher Scientific) mass spectrometer. The heated electrospray ionisation (HESI) source was operated at 375 °C and 3.5 kV in positive ion mode. The capillary temperature was 380°C; sheath and auxillary gases were 60 and 20 arbitrary units, respectively. Spectra were obtained over range of 200-2000 m/z at 60,000 mass resolution. Data were converted to mzXML using ProteoWizard. Features were identified using xcms[22] with 10 parts-per-million (ppm) spectral binning. Peak areas were normalised to the relevant internal standard peak area and protein content. Lipid identification was performed by mass matching to an in-house database and LIPID MAPS® structure database[23]. Multivariate statistics were performed using Simca 17 (Sartorius, Gottingen, Germany) to conduct principal components analysis (PCA) with Pareto scaling.

**Statistical Analysis**

Graphs and heatmaps were prepared using GraphPad prism software (version 10.5.0). Data is presented as mean ± standard error of the mean (SEM). P-values of <0.05 were considered statistically significant and presented on graphs as * p<0.05, ** p<0.01, *** p<0.001 and p<0.0001. P-values were calculated using GraphPad prism software (version 11.0.2) using either a One Way ANOVA with Dunnett's multiple comparison test, Two Way ANOVA with Tukey's multiple comparisons test or Paired T-tests as appropriate.
